# Loss of heterozygosity of essential genes represents a widespread class of potential cancer vulnerabilities

**DOI:** 10.1101/534529

**Authors:** Caitlin A. Nichols, William J. Gibson, Meredith S. Brown, Jack A. Kosmicki, John P. Busanovich, Hope Wei, Laura M. Urbanski, Naomi Curimjee, Ashton C. Berger, Galen F. Gao, Andrew D. Cherniack, Sirano Dhe-Paganon, Brenton R. Paolella, Rameen Beroukhim

## Abstract

Alterations in non-driver genes represent an emerging class of potential therapeutic targets in cancer. Hundreds to thousands of non-driver genes undergo loss of heterozygosity (LOH) events per tumor, generating discrete differences between tumor and normal cells. Here we interrogate LOH of polymorphisms in essential genes as a novel class of therapeutic targets. We hypothesized that monoallelic inactivation of the allele retained in tumors can selectively kill cancer cells but not somatic cells, which retain both alleles. We identified 5664 variants in 1278 essential genes that undergo LOH in cancer and evaluated the potential for each to be targeted using allele-specific gene-editing, RNAi, or small-molecule approaches. We further show that allele-specific inactivation of either of two essential genes (*PRIM1* and *EXOSC8*) reduces growth of cells harboring that allele, while cells harboring the non-targeted allele remain intact. We conclude that LOH of essential genes represents a rich class of non-driver cancer vulnerabilities.

## Introduction

Despite progress in precision cancer drug discovery, few highly selective therapies exist in the clinic. A current paradigm focuses on drugging driver alterations in cancer; however, many driver genes have proven difficult to target therapeutically^1–3^, and in many cancers no easily targeted drivers exist. Alterations in non-driver genes represent an alternative target class that merits further investigation.

Loss of heterozygosity (LOH) may generate cancer-specific vulnerabilities by eliminating genetic redundancy in cancer cells. LOH occurs when a cancer cell that is originally heterozygous at a locus loses one of its two alleles at that locus, either by simple deletion of one allele (copy-loss LOH), or by deletion of one allele accompanied by duplication of the remaining allele (copy-neutral LOH). In either case, the cancer cell then relies on the gene products encoded by a single allele, in contrast to normal cells, which retain both alleles. When a cancer cell undergoes LOH of an essential gene, further loss or inhibition specifically of the allele retained in the tumor should not be tolerated, whereas normal cells will be able to survive relying solely on the remaining allele^4^ (Figure 1A). We term this target class GEMINI vulnerabilities, after the twins from Greek mythology Castor and Pollux, one of which was mortal and the other immortal.

**Figure 1:**
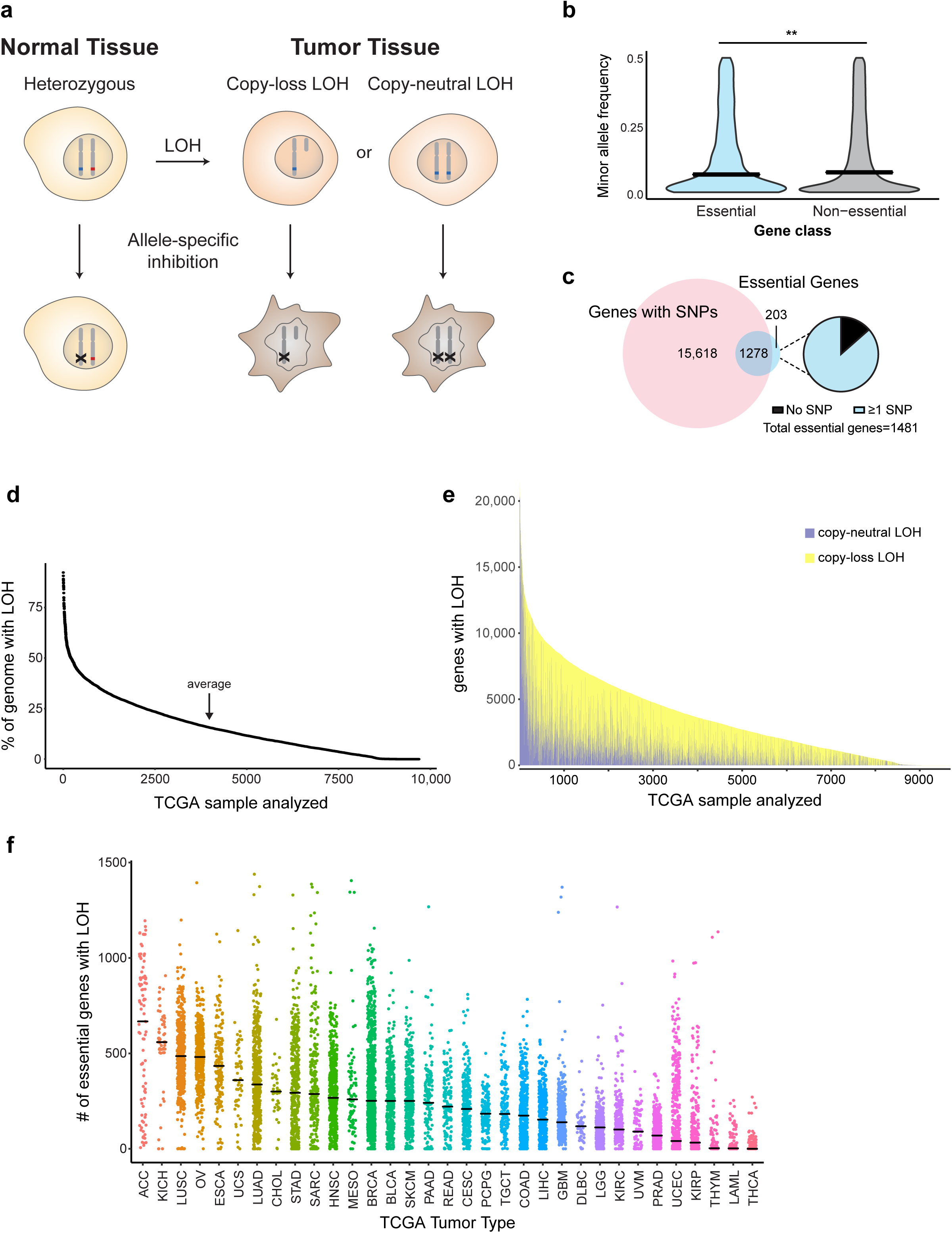
Genomic rates of LOH and allelic variation in normal and cancer genomes. **a.** Schematic indicating how loss of heterozygosity (LOH) of essential genes represents a potentially targetable difference between cancer and normal cells. **b.** Violin plot of minor allele frequency of polymorphisms in essential versus non-essential genes in the ExAC cohort. Intersecting lines represent median values: essential = 0.141, non-essential = 0.146; one-tailed Student’s t-test, **p < 0.01. **c.** (Left) Overlap between genes with common polymorphisms in the ExAC database (pink circle) and essential genes (blue circle). (Right) Fraction of essential genes with common polymorphisms. **d.** Percent of genome affected by LOH across 9,686 cancers from TCGA. **e.** Stacked histogram representing the number of genes with copy-loss (yellow) or copy-neutral LOH (purple) across 9,686 cancers from TCGA. **f.** Dot plot of the number of essential genes affected by LOH across 33 TCGA tumor types. Tumor types are indicated by TCGA abbreviations (see https://gdc.cancer.gov/resources-tcga-users/tcga-code-tables/tcga-study-abbreviations). Each dot represents an individual sample. Lines indicate median values.

While previous reports have described individual GEMINI vulnerabilities^5,6^, these studies have not systematically evaluated the landscape of potential targets, taking into account genome-scale assessments of gene essentiality, variation in human genomes, and rates of LOH across cancers. Open questions include which essential genes exhibit widespread variation in human populations and frequent LOH in cancers, providing potential GEMINI vulnerabilities, and at what rates these vulnerabilities occur. Moreover, different GEMINI vulnerabilities may require different therapeutic approaches to exploit them, due to the location of the variant within each gene and its effects on the amino acid composition of the protein. These differences have not been explored. Furthermore, GEMINI vulnerabilities have never been validated in isogenic systems to confirm specificity.

To address these questions, we integrated genome-scale copy number, germline allelic variation, and gene essentiality data to identify a list of polymorphisms in cell essential genes that undergo LOH in cancer, serving as a compendium of potential GEMINI targets. We also performed proof-of-principle validation of GEMINI vulnerabilities for two candidate genes in this list, *PRIM1* and *EXOSC8*, using allele-specific CRISPR in both patient-derived and isogenic models. These results rigorously validate the GEMINI class of vulnerabilities and define its potential scope.

## Results

### Integration of genome-scale loss of heterozygosity and germline variation analyses reveals a class of frequent cancer-specific alterations

To identify potential targets for our approach, we first characterized the landscape of cell-essential genes. We integrated genome-wide gene essentiality data from loss-of-function genetic screens and CCLE cell lines to conservatively estimate 1481 cell-essential genes (Supplementary Table 1; Methods). This list is enriched for genes involved in essential cellular processes including rRNA processing, mRNA splicing, and translation (functional enrichment analysis performed with DAVID^7,8^, version 6.8; Supplementary Table 2).

We then assessed germline heterozygosity resulting from normal human genetic variation in coding regions and 5’ and 3’ untranslated regions (UTRs) using allele frequencies across 60,706 individuals in the Exome Aggregation Consortium database^9^. Variants at 90,409 loci were observed to be present among at least 1% of alleles. As expected, polymorphisms in essential genes are slightly less common than those in non-essential genes (median minor allele frequency: essential = 0.141, non-essential = 0.146; p = 0.005, one-tailed Student’s t-test; Figure 1B). However, essential genes still contain an abundance of genetic variation: 86% (1278/1481) harbor at least one common germline variant (Figure 1C), with 49% (730/1481) harboring at least one missense variant. The median essential gene contains 3 germline polymorphisms. The median polymorphism in an essential gene is heterozygous in 13.9% of individuals (Supplementary Figure 1A).

We were interested in how much of this heterozygosity in essential genes is lost in cancer. Loss of heterozygosity (LOH) in cancer frequently results from copy number alterations (CNAs) that can alter dozens to thousands of genes in cancer genomes^10,11^. Most LOH is due to strict copy-loss (copy-loss LOH), where allelic loss occurs in the context of a decrease in gene copy number.

However, copy-neutral LOH is also frequently observed, whereby an allele is lost but the number of gene copies remains the same due to a duplication event, or in some cases, even increases. LOH has been frequently described^11,12^, but to our knowledge there has not yet been a systematic analysis of the frequency of LOH events across cancer types.

We therefore analyzed copy number and LOH calls from 9,686 patient samples across 33 TCGA tumor types (Methods). On average and across all cancers, 16% of genes undergo LOH (Figure 1D). Genome-wide LOH rates vary widely by tumor type, however, ranging from a median of 45% in adenoid cystic carcinoma to 0.01% in thyroid carcinoma (Supplementary Figure 1B). Approximately 28% of genes undergoing LOH undergo copy-neutral LOH (Figure 1E), and on average across all cancers, 4.4% of all genes undergo copy-neutral LOH.

Rates of LOH are no lower for cell-essential genes relative to the rest of the genome (essential: 16.4%, non-essential: 15.6%; p=1, one-tailed Student’s t-test; Supplementary Figure 1C), suggesting that LOH of essential genes does not impose negative selection pressure. As a result, tumors harbored an average of 189 essential genes with LOH (Figure 1F).

We hypothesized that the widespread nature of LOH of essential genes could represent a new opportunity to target essential genes that are heterozygous in normal tissue but undergo LOH in cancer. Among individuals with heterozygous SNPs within an essential gene, cancer cells with LOH of that gene would rely solely on the gene product encoded by one allele, in contrast to somatic cells, which would retain both alleles. We therefore hypothesized that allele-specific inactivation of the allele that had been retained in the cancer would selectively kill the cancer cells (Figure 1A).

Our analysis identified 5664 polymorphisms in 1278 cell-essential genes, representing a compendium of potential GEMINI vulnerabilities (Supplementary Table 3). These GEMINI genes are enriched for similar pathways as the wider set of essential genes, including rRNA processing, mRNA splicing, and translation (functional enrichment analysis performed with DAVID^7,8^, version 6.8; Supplementary Table 4). Among the 5664 GEMINI variants, 1688 lead to missense changes in amino acid composition of an essential protein, raising the possibility that they could be distinguished by molecules that interact with the protein directly. We focused on two of these missense SNPs for further functional analysis.

### Validation of *PRIM1*^rs2277339^ as a GEMINI vulnerability

Variants residing in putative CRISPR protospacer adjacent motif (PAM) sites have previously been shown to enable allele-specific gene disruption^13–15^. For nuclease activity, *S. pyogenes* Cas9 requires a PAM site of the canonical motif 5’-NGG-3’ downstream of the 20-nucleotide target site; deviations from this motif abrogate Cas9-mediated target cleavage^16,17^. Therefore, we hypothesized that in the case in which one allele of a SNP generates a novel PAM site, Cas9 would be able to disrupt the “CRISPR-sensitive” (S), G allele that maintains the PAM sequence while leaving the other, “CRISPR-resistant” (R) allele intact (Figure 2A).

**Figure 2:**
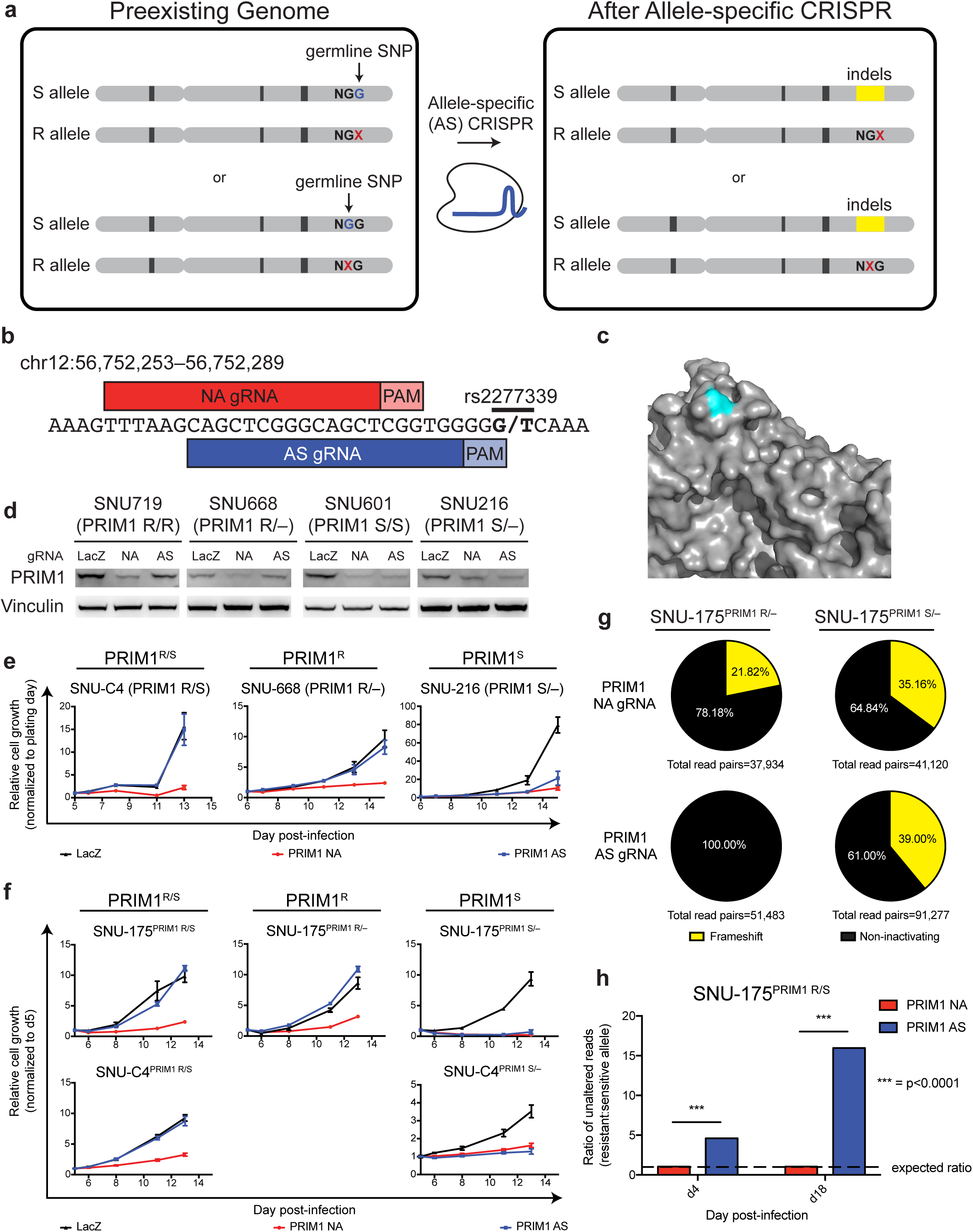
Validation of *PRIM1*^rs2277339^ as a GEMINI vulnerability. **a.** Schematic indicating allele-specific CRISPR approach. “Preexisting genome” represents individuals heterozygous for a germline SNP in a *S. pyogenes* Cas9 protospacer adjacent motif (PAM) site. A “G” allele (blue) in the PAM retains Cas9 activity at the target site, making this allele CRISPR-sensitive (S). An allele other than “G,” represented by “X” (red) abrogates Cas9 activity at the target site, making this allele CRISPR-resistant (R). Expression of an allele-specific (AS) CRISPR sgRNA targeting the polymorphic PAM site leads to specific inactivation of the S allele. **b.** Schematic of *PRIM1* SNP rs2277339 locus showing target sites for positive control, non-allele specific (NA) sgRNA and experimental, allele-specific (AS) sgRNA. Alleles appear in bold. **c.** Crystal structure of *PRIM1* gene product^92^ shows the amino acid encoded by rs2277339 (teal) lies on the surface of the primase catalytic subunit (grey) near a potentially small-molecule accessible location. **d.** Immunoblot of PRIM1 protein levels in indicated patient-derived cell lines expressing LacZ, PRIM1 NA, or PRIM1 AS sgRNA. **e.** Representative growth curves of indicated patient-derived cell lines expressing LacZ (black), PRIM1 NA (red), or PRIM1 AS (blue) sgRNA, as measured by CellTiter-Glo luminescence, relative to day of assay plating. n = 5 technical replicates; error bars represent s.d. See Supplementary Figure 2 for additional biological replicates. **f.** Representative growth curves of indicated isogenic cell lines expressing LacZ (black), PRIM1 NA (red), or PRIM1 AS (blue) sgRNA, as measured by CellTiter-Glo luminescence, relative to day of assay plating. n = 5 technical replicates; error bars represent s.d. See Supplementary Figure 2 for additional biological replicates. **g.** Disruption of *PRIM1* in isogenic hemizygous *PRIM1* resistant (PRIM1^R^) or *PRIM1* sensitive (PRIM1^S^) cells expressing PRIM1 NA or AS sgRNA. Non-inactivated alleles (representing unaltered alleles or alleles with in-frame insertions or deletions, black) and alleles with frameshift alterations (yellow) were assessed by deep sequencing of *PRIM1* four days post-infection with sgRNA. **g.** Ratio of unaltered resistant to unaltered sensitive alleles of *PRIM1*^rs2277339^ in *PRIM1* heterozygous cells (PRIM1^R/S^) at day 4 and day 18 post-infection with NA or AS sgRNA as assessed by deep sequencing. Dashed line indicates expected ratio of unaltered R: S alleles if an sgRNA targets either allele with equal frequency. Chi-square with Yates correction, ***p<0.0001.

We identified such a SNP in the essential gene *PRIM1* as a promising candidate for proof-of-principle validation. *PRIM1* encodes the catalytic subunit of DNA primase and has previously been determined to be an essential gene^18–20^. It contains two common SNPs, of which one (rs2277339) leads to a change in the amino acid sequence: a T to G substitution resulting in conversion of an aspartate on the protein surface to an alanine (Figure 2B and 2C; Supplementary Figure 2A). The minor allele is common (minor allele frequency = 0.177), leading to heterozygosity at this locus in 29% of individuals represented in the ExAC database. This locus also undergoes frequent LOH. Across the 33 cancer types profiled, LOH was observed at the rs2277339 locus in 9% of cancers, including 21% of lung adenocarcinomas, 18% of ovarian cancers, and 17% of pancreatic cancers (Supplementary Figure 2B).

*PRIM1*^rs2277339^ lies in a polymorphic PAM site—the “CRISPR-sensitive,” G allele generates a canonical *S. pyogenes* Cas9 PAM site, while the “CRISPR-resistant,” T allele disrupts the NGG PAM motif. We tested allele-specific *PRIM1* disruption using an allele-specific (AS) CRISPR single guide RNA (sgRNA) designed to target only the G allele at rs2277339, encoding the alanine version of the protein (Figure 2B). In the context of CRISPR experiments, because the G allele should be *sensitive* to allele-specific disruption, we use an “S” to designate cells with this allele and an “R” to designate cells with the other, *resistant* allele: for example, PRIM1^S/–^ and PRIM1^R/S^ genotypes reflect cells with one copy of the sensitive G allele and cells with one copy of each allele, respectively.

We transduced four patient-derived cancer cell lines that naturally exhibit either allele with AS sgRNA and verified that AS sgRNA disrupts *PRIM1* in PRIM1^S^ genetic contexts (Figure 2D). PRIM1^S/–^ and PRIM1^S/S^ cells expressing AS sgRNA show decreased proliferation relative to LacZ-targeting control, whereas cells retaining the resistant allele (PRIM1^R/–^, PRIM1^R/R^, or PRIM1^R/S^) show no such defects (Figure 2E, Supplementary Figure 2C–F).

The specificity of the AS sgRNA for PRIM1^S^ cell lines was not due to a lack of Cas9 activity or *PRIM1* essentiality in the PRIM1^R^ cell lines. We confirmed this finding by transducing four cell lines with a non-allele specific (NA) *PRIM1*-targeting sgRNA. We successfully ablated PRIM1 expression in all contexts (Figure 2D), and cells expressing *PRIM1*-targeting sgRNA showed dramatic and significant decreases in proliferation relative to *LacZ*-targeting control even in cases where expression of the AS sgRNA did not significantly limit growth (p<0.01 in all cases, one-tailed Student’s t-test; Figure 2E, Supplementary Figure 2C–F).

We further tested isogenic cell lines harboring either allele. Using SNU-175 and SNU-C4 cells, which are heterozygous for *PRIM1*^rs2277339^, as a base, we transiently transfected a vector expressing Cas9 and two sgRNAs that flank the *PRIM1* gene. We then screened single cell clones for *PRIM1* deletion by PCR. Among deletion-positive clones, we identified heterozygous (PRIM1^R/S^), hemizygous sensitive (PRIM1^S^), and hemizygous resistant (PRIM1^R^) lines (Supplementary Figure 3A–E). Using these isogenic cells, we confirmed PRIM1^S/–^ cells expressing AS sgRNA show decreased proliferation relative to LacZ-targeting control, whereas cells retaining the resistant allele (PRIM1^R/–^ or PRIM1^R/S^) show no such defects (Figure 2F, Supplementary Figure 2G–K).

**Figure 3:**
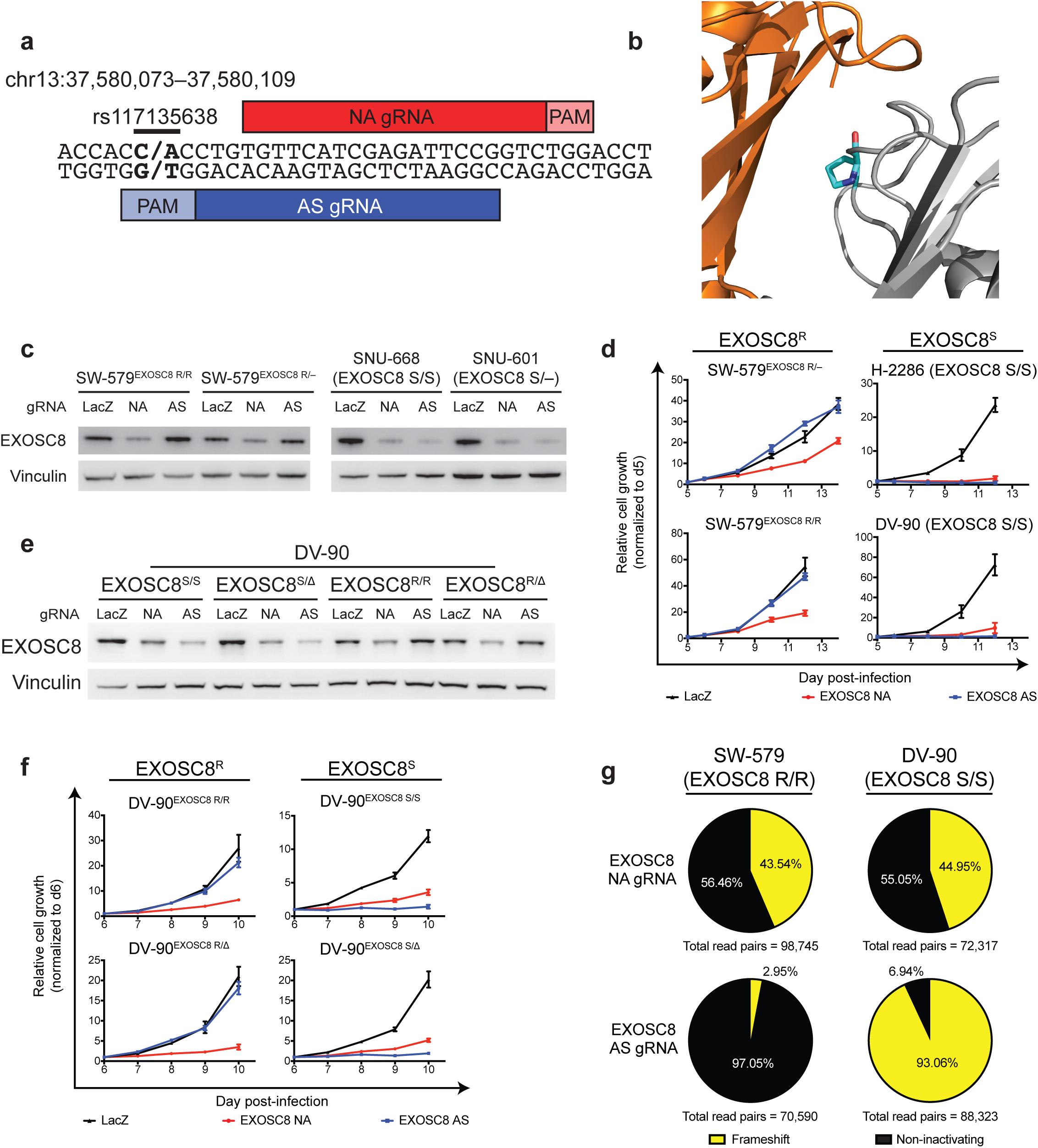
Validation of *EXOSC8*^rs117135638^ as a GEMINI vulnerability. **a.** Schematic of *EXOSC8* SNP rs117135638 locus showing target sites for positive control, non-allele specific (NA) sgRNA and experimental, allele-specific (AS) sgRNA. Alleles appear in bold. **b.** Crystal structure of *EXOSC8* gene product, Rrp43^93^ (grey) shows the amino acid encoded by rs117135638 (teal) lies on the surface of the Rrp43 protein near the interface with exosome complex subunit Mtr3 (pink). **c.** Immunoblot of EXOSC8 protein levels in indicated patient-derived and isogenic cell lines expressing LacZ, EXOSC8 NA, or EXOSC8 AS sgRNA. **d.** Representative growth curves of indicated patient-derived and isogenic cell lines expressing LacZ (black), EXOSC8 NA (red), or EXOSC8 AS (blue) sgRNA, as measured by CellTiter-Glo luminescence, relative to day of assay plating. n = 5 technical replicates; error bars represent s.d. See Supplementary Figure 4 for additional biological replicates. **e.** Immunoblot of EXOSC8 protein levels in indicated isogenic cell lines expressing LacZ, EXOSC8 NA, or EXOSC8 AS sgRNA. **f.** Representative growth curves of indicated isogenic cell lines expressing LacZ (black), PRIM1 NA (red), or PRIM1 AS (blue) sgRNA, as measured by CellTiter-Glo luminescence, relative to day of assay plating. n = 5 technical replicates; error bars represent s.d. See Supplementary Figure 4 for additional biological replicates. **g.** Disruption of *EXOSC8* in patient-derived *EXOSC8* resistant (EXOSC8^R^) or *EXOSC8* sensitive (EXOSC8^S^) cells expressing EXOSC8 non-allele specific (NA) positive control sgRNA or allele-specific (AS) experimental sgRNA. Non-inactivated alleles (representing unaltered alleles or alleles with in-frame insertions or deletions, black) and alleles with frameshift alterations (red) were assessed by deep sequencing of *EXOSC8* four days post-infection with sgRNA.

Within these isogenic lines, we also confirmed that AS CRISPR disrupts *PRIM1* in a *PRIM1*^rs2277339^-dependent manner using deep sequencing. Four days post-infection with the NA sgRNA, isogenic PRIM1^S^ and PRIM1^R^ cells showed comparable fractions of disrupted alleles (Figure 2G), suggesting both lines exhibit similar levels of Cas9 activity. However, while PRIM1^S^ cells expressing AS sgRNA showed nearly 40% disrupted alleles, resistant cells under the same condition showed 0 disrupted alleles (p<0.0001, Chi-square with Yates correction; Figure 2G). This result confirms that AS PRIM1 sgRNA targets PRIM1 in a SNP-specific manner.

We also verified that allele-specific inactivation of essential genes is possible in a heterozygous context. Gene-disrupting indels introduced by the error-prone non-homologous end joining (NHEJ) repair pathway make distinguishing the original genotype (S or R) of an edited allele challenging. Therefore, we compared the number of unaltered reads of each allele in heterozygous cells expressing either NA or AS sgRNA. If the sgRNA disrupts *PRIM1* in a non-allele specific fashion, we would expect a ratio of 1 between unaltered reads of each allele. We infected a Cas9-stable PRIM1^R/S^ line with NA or AS sgRNA as described above and sequenced the target loci 4 and 18 days post-infection. As expected, we saw no substantial difference in the number of unaltered reads between the two alleles in PRIM1^R/S^ cells expressing NA sgRNA (Figure 2H). In contrast, PRIM1^R/S^ cells expressing AS sgRNA showed significantly more unaltered reads from the resistant allele compared to the sensitive allele, with a ratio that increased over time (p<0.0001, Chi-square with Yates correction). We conclude that the AS sgRNA disrupts *PRIM1* in a SNP-specific manner even in a heterozygous context.

### Validation of *EXOSC8*^rs117135638^ as a GEMINI vulnerability

We also performed proof-of-principle validation for another candidate SNP in the essential gene *EXOSC8*. *EXOSC8* codes for Rrp43, a component of the RNA exosome. The RNA exosome is an essential multi-protein complex involved in RNA degradation and processing, including processing of pre-rRNA^21–23^. Two common SNPs have been described within *EXOSC8*, one of which (rs117135638) represents a C to A change in DNA sequence; this SNP leads to a proline to histidine substitution on the interface between Rrp43 and exosome complex member Mtr3 (Figure 3A and 3B; Supplementary Figure 4A). This candidate SNP is heterozygous in 2% of individuals and undergoes LOH in 29% of cancers, including 72% of lung squamous cell carcinomas, 62% of ovarian cancers, 46% of lung adenocarcinomas, and 40% of breast cancers (Supplementary Figure 4B).

**Figure 4:**
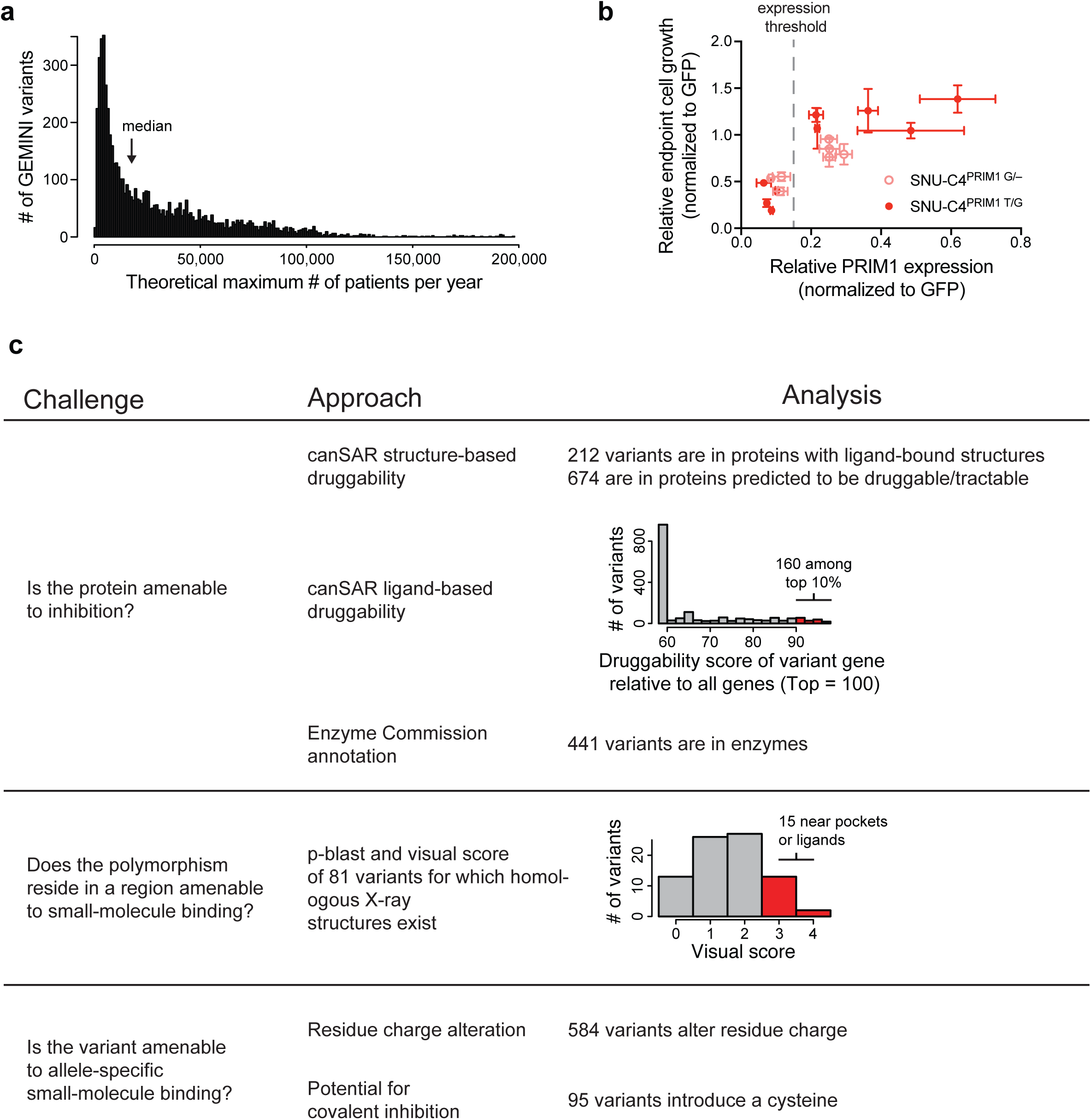
Potential therapeutic approaches to targeting GEMINI vulnerabilities. **a.** Number of GEMINI variants (vertical axis) plotted against the number of patients per year whose tumors might respond to therapeutics targeting those variants (i.e., have lost the resistant allele from a heterozygous germline; horizontal axis). Bin width = 1000 patients. **b.** Growth of heterozygous and hemizygous cells expressing positive control, non-allele specific *PRIM1*-targeting shRNAs versus *PRIM1* mRNA expression. Cell growth measured by CellTiter-Glo luminescence relative to day 2 post-infection and shGFP (n = 5 technical replicates). *PRIM1* mRNA expression assessed by qRT-PCR (n = 3 technical replicates). Dashed grey line indicates *PRIM1* expression threshold below which substantial decreases in cell viability are observed. **c.** Summary table representing challenges to developing allele-specific small molecules that target GEMINI vulnerabilities and associated analyses to prioritize targets.

We first tested allele-specific *EXOSC8* disruption using allele-specific (AS) RNA sgRNAs designed to target only the C allele at rs117135638, encoding the proline version of the protein. We designated cells as EXOSC8^S^ (for “sensitive”) if they contained this allele, and as EXOSC8^R^ (for “resistant”) if they contained the A allele. We transduced four patient-derived cancer cell lines with AS sgRNA and verified that AS sgRNA disrupts EXOSC8 protein expression in EXOSC8^S^ genetic contexts (Figure 3C). EXOSC8^S/S^ and EXOSC8^S/–^ cells expressing AS sgRNA showed decreased proliferation relative to LacZ-targeting control, whereas cells retaining the resistant allele (EXOSC8^R/–^ or EXOSC8^R/R^) showed no such defects (Figure 3D, Supplementary Figure 4C–F).

We next verified that the specificity of the AS sgRNA for EXOSC8^S^ cell lines was not due to a lack of Cas9 activity or *EXOSC8* essentiality in the EXOSC8^R^ cell lines. We transduced all four cell lines with a non-allele specific (NA) targeting sgRNA, which successfully ablated EXOSC8 expression in all contexts (Figure 3C). Cells expressing *EXOSC8*-targeting sgRNA showed significant decreases in proliferation relative to *LacZ*-targeting control (p<0.01 in all cases, one-tailed Student’s t-test), confirming that this gene is cell-essential (Figure 3D, Supplementary Figure 4C-F).

We also determined that both copy-loss and copy-neutral LOH of *EXOSC8* represents a vulnerability in an isogenic context. We generated diploid and single-copy knockout isogenic cells representing EXOSC8^S^ and EXOSC8^R^ genotypes by Cas9-mediated homology-directed repair (HDR) editing (Methods; Supplementary Figure 3F–I), and then infected these isogenic Cas9-stable lines with constructs expressing EXOSC8 NA or AS sgRNA. As expected, EXOSC8 NA sgRNA ablated EXOSC8 expression in all contexts, while AS sgRNA ablated EXOSC8 expression only in EXOSC8^S^ cells (Figure 3E). EXOSC8^S/S^ and EXOSC8^S/Δ^ cells expressing AS sgRNA showed decreased proliferation relative to *LacZ*-targeting control, whereas cells retaining the resistant allele (EXOSC8^R/R^ and EXOSC8^R/Δ^) showed no such defects (Figure 3F, Supplementary Figure 4G– J).

We also confirmed that AS CRISPR disrupts *EXOSC8* in a SNP-dependent manner using deep sequencing. Four days post-infection with the NA sgRNA, SW-579 (EXOSC8 R/R) and DV-90 (EXOSC8 S/S) cells showed approximately equal fractions of disrupted alleles (Figure 3G). However, while DV-90 sensitive cells expressing AS sgRNA showed upwards of 90% disrupted alleles, SW-579 resistant cells under the same condition showed fewer than 3% disrupted alleles, a significant difference (p<0.0001, Chi-square with Yates correction). This result confirms that EXOSC8 AS sgRNA targets *EXOSC8* in a SNP-specific manner.

### Potential Approaches to Targeting GEMINI Vulnerabilities in Humans

We were interested in understanding the potential scope of patients that could benefit from therapeutic approaches targeting GEMINI vulnerabilities. For each GEMINI variant, we calculated the number of new patients per year that exhibit LOH of the hypothetical “targetable” allele (Methods). Across the 33 tumor types we profiled, the median GEMINI vulnerability could be targetable in 17,747 patients per year (Figure 4A). *PRIM* ^rs2277339^ and *EXOSC8*^rs117135638^ could be targetable in a theoretical 22,470 and 5,307 patients per year, respectively (Supplementary Figure 2A and 4A).

The major challenge to exploiting GEMINI vulnerabilities is identifying means to target them in humans. Three approaches that may be contemplated are DNA-targeting CRISPR effectors (e.g., Cas9), RNA-targeting approaches (e.g., RNAi), and allele-specific small molecules. We characterized each GEMINI vulnerability according to criteria that would indicate its amenability to targeting by each of these approaches.

First, we analyzed the list of GEMINI vulnerabilities to identify polymorphisms whose targeting on the DNA level may enable allele-selective gene disruption by a CRISPR-based approach. For this analysis, we included both the canonical *S. pyogenes* PAM, NGG, as well as the weaker, non-canonical PAM, NAG^17,24^. Of the 4648 GEMINI vulnerabilities in open reading frames, 23% (1088/4648, or 19% of all GEMIMI vulnerabilities) generate a PAM site in one allele but not the other, suggesting the potential for allele-specific knockout (Supplementary Table 3).

In theory, every GEMINI variant could be the target of allele-specific RNAi reagents. However, it is possible that, for some GEMINI variants, RNAi reagents would be unable to suppress expression sufficiently to reduce cell viability, or that sufficient allelic specificity might not be achieved. For example, we tested the hypothesis that *PRIM1*^rs2277339^ may be targetable in an allele-specific manner using RNAi. For these experiments, we refer to the *PRIM1* alleles by their identifying nucleotide; for example, heterozygous cells are referred to as PRIM1^T/G^, and cells hemizygous for the minor allele are referred to as PRIM1^G/–^. We first sought to determine the level of PRIM1 knockdown required to substantially decrease cell proliferation. Accordingly, we infected hemizygous and heterozygous isogenic cells with non-allele specific *PRIM1*-targeting shRNAs and assessed *PRIM1* expression and cell growth. We observed that substantial decreases in cell growth were possible under conditions of robust *PRIM1* knockdown (>85%) (Figure 4B).

We then asked whether allele-specific shRNAs targeting the *PRIM1*^rs2277339^ locus could decrease growth in cells representing the fully matched genotype. PRIM1^T/–^ and PRIM1^G/–^ cells were infected with constructs encoding fully complementary shRNAs tiling across the SNP and assessed for growth. Only one shRNA, shG7 (targeting the minor, G allele at position 7) significantly reduced cell growth relative to GFP-targeting control (Supplementary Figure 5A–B). We then selected the four *PRIM1* SNP-targeting shRNAs that yielded the lowest average cell growth relative to GFP-targeting control and assessed their ability to decrease cell growth in an allele-specific manner. Heterozygous cells (PRIM1^T/G)^ and hemizygous cells of the targeted genotype (PRIM1^T/–^ or PRIM1^G/–^) were infected with constructs encoding the appropriate shRNA. No putative allele-specific shRNAs were found to significantly decrease cell growth in hemizygous cells of the targeted genotype relative to heterozygous cells (Supplementary Figure 5C–D). We conclude that *PRIM1*^rs2277339^ may not represent an optimal candidate for allele-specific shRNA-mediated inhibition.

Given the large number of additional GEMINI variants that may be suitable for RNAi-mediated targeting, we sought to prioritize GEMINI genes that may be amenable to allele-specific inhibition using mRNA-targeting approaches. RNAi-mediated knockdown of some essential genes may be more effective at inducing cell death than others, based on differential expression thresholds required for cell survival^25–27^. We hypothesized that GEMINI genes representing strong dependencies in shRNA screens would be most amenable to potential targeting using an RNAi-based therapeutic. We therefore analyzed shRNA data representing 17,212 genes in 712 cell lines^28^, including 1183 GEMINI genes, and looked for genes whose suppression led to at least a moderately strong response in most of the cell lines (median DEMETER2 score < −0.5; Methods). Approximately 35% of GEMINI genes (413/1183), including PRIM1 (median DEMETER2 score = −0.52), fit this category, representing 35% (1804/5196) of GEMINI vulnerabilities (Supplementary Table 3). In comparison, only 3.6% of all genes profiled (623/17,212) passed this dependency threshold, indicating a significant enrichment for GEMINI genes (p < 0.0001, binomial proportion test). However, this level of dependency was not observed for all GEMINI or essential genes. For example, the median DEMETER2 score for *EXOSC8* was only −0.14, despite our and others’ extensive data showing its essentiality in multiple cell types^21^. These results raise the possibility that RNAi-based approaches may not be able to exploit many GEMINI vulnerabilities.

Both CRISPR- and RNAi-based therapeutic approaches suffer from difficulties in effectively delivering reagents to all cancer cells in an animal. Small molecule–based approaches can overcome such delivery issues, but substantial obstacles exist to developing allele-specific small molecules that target GEMINI vulnerabilities. These challenges include identifying GEMINI genes that are amenable to small-molecule inhibition, determining which GEMINI variants lie near potentially druggable pockets, and predicting which GEMINI variants are most likely to facilitate allele-specific drug binding. We therefore analyzed 1749 protein-altering GEMINI vulnerabilities (missense, insertion, and deletion variants) to prioritize candidates for potential allele-specific drug development (Supplementary Table 5).

To identify GEMINI genes that may be amenable to small-molecule inhibition, we annotated those containing protein-altering alleles using the canSAR Protein Annotation Tool^29,30^ (cansar.icr.ac.uk; Methods). This tool uses publicly available structural and chemical information to generate structural and ligand-based druggability scores. While these scores do not necessarily reflect potential for allele-specific small molecule inhibition based on the GEMINI variant of interest, they may nonetheless allow prioritization of targets based on general druggability. This analysis found that of the 1734 protein-altering variants in genes assessed by canSAR, 12% (212) reside in proteins with a small molecule ligand–bound structure (Figure 4C). Additional assessments of potential small-molecule binding sites on structures with and without existing ligands indicated that 39% of protein-altering variants (674) lie in proteins with molecular structures that are predicted to be druggable (drug-like compound modulates activity in vivo) or tractable (tool compound modulates activity in vitro) (Figure 4C). Furthermore, 160 GEMINI variants reside in proteins in the top 90th percentile of ligand-based druggability as assessed by the physiochemical properties of small molecules tested against the protein or its homologs (Figure 4C). We also found that 25% of protein-altering GEMINI variants (441/1734) reside in enzymes as defined by their annotation with an Enzyme Commission (EC) number^31^ (Figure 4C).

To assess which variants may reside in protein regions amenable to small molecule binding, we performed a p-blast of the 1749 protein-altering variants against protein sequences for molecular structures found in the Protein Data Bank^32^ (rcsb.org; Methods). This analysis identified 153 variants characterized in a homologous structure. We then visually scored 81 missense and indel variants in X-ray crystal structures for their proximity to solvent-exposed pockets or known small-molecule binding sites using a scale of 0 to 4 (Methods). Of the variants analyzed, 15 were near a potential binding pocket on the surface of the protein (score = 3), with two of these pockets containing a small molecule ligand (score = 4) (Figure 4C).

We also assessed protein-altering GEMINI variants to prioritize those that may be most amenable for allele-specific small-molecule inhibition. For this analysis, we scored variants that altered the number or sign of residue charges. For example, a variant that changes the charge of a residue from neutral to negative or that adds an additional negative charge through an inserted residue would qualify as a charge-altering variant. Of the 1749 protein-altering variants, 584 induced a change in residue charge (Figure 4C). We further hypothesized that variants introducing a cysteine residue could provide additional allele selectivity by enabling the potential development of a covalent inhibitor. Among the missense and indel GEMINI variants, 95 generate a cysteine in one allele.

We then integrated each of these analyses to characterize the potential druggability landscape of these protein-altering GEMINI vulnerabilities. Every variant was given a score from 0 to 7 based on the number of analyses in which it scored among the top candidates. One variant, *TGS1*^rs7823773^, earned a score of 6, including in the visual scoring and cysteine categories. Nine additional variants earned a score of 5 (Supplementary Table 5). These may be among the highest-priority candidates for further exploration.

## Discussion

Leveraging synthetic lethal interactions in cancer cells represents a promising avenue to targeting genomic differences between tumor and normal tissue. Synthetic lethality between genes occurs when singly inactivating one gene or the other maintains viability, but inactivating both genes simultaneously causes lethality^33^. Over the past 20 years, many efforts have been directed toward discovering synthetic lethal interactions with genetic driver alterations of oncogenes and tumor suppressor genes^34,35^. However, the number of genetically activated oncogenes and inactivated tumor suppressor genes in any given tumor is limited and, in many cancer types, is vastly outnumbered by genetically altered non-driver genes (e.g., due to passenger events). Therefore, identifying synthetic lethalities with genetic alterations affecting non-driver genes (also termed “collateral lethalities”^36^) would increase the scope of potential therapeutic approaches. While individual GEMINI genes have been described previously^4^, our work integrated genome-wide assessments of gene essentiality, genetic variation, and LOH to generate the first systematic analysis of this target class.

GEMINI vulnerabilities represent one of four classes of collateral lethalities. In addition to GEMINI vulnerabilities, deletion of paralogs can result in dependency on the remaining paralog; loss or gain of function of a non-driver pathway can lead to dependencies on alternative non-driver pathways^36^; and hemizygous loss of essential genes can result in dependency on the remaining copy (CYCLOPS)^25,26^.

Prior analyses have indicated CYCLOPS genes to be the most frequent class of these synthetic lethal interactions^26,27^, but we find that GEMINI vulnerabilities provide similar numbers of potential targets. In comparison, fewer paralog dependencies have been described^27,37–40^. Larger numbers of paralog vulnerabilities have been predicted^41^, but it is unclear whether these predictions represent viable candidates^36^. The 1278 GEMINI genes that we identified also exceed the 299 known driver genes^42^, many of which are proposed therapeutic targets. Expanding the search for GEMINI vulnerabilities beyond pan-essential genes to include variants in lineage-specific essential genes may also increase the number of potential GEMINI targets.

In comparison to CYCLOPS, targeting GEMINI vulnerabilities has two distinct advantages. First, whereas CYCLOPS genes must lie in regions of copy loss, GEMINI genes encompass genes that undergo both copy-loss and copy-neutral LOH. Second, while CYCLOPS vulnerabilities rely on relative differences between tumor and normal cells (differential expression of target genes), GEMINI vulnerabilities exploit absolute differences (the presence or absence of the allele that has undergone LOH). Thus, the possibility of allele-specific targeting presented by GEMINI genes may widen prospective therapeutic windows. In 269 cases, GEMINI vulnerabilities we detected reside in CYCLOPS genes^26^ (Supplementary Table 3). If GEMINI and CYCLOPS vulnerabilities are additive, targeting these genes might offer an even wider therapeutic window in cancers where CYCLOPS-GEMINI genes suffer LOH due to copy loss. However, individual GEMINI alterations may be less common among patients than individual CYCLOPs alterations due to the requirement that the germline genome be heterozygous at the GEMINI locus.

Like any target class, we expect resistance mechanisms to arise in response to targeting GEMINI vulnerabilities. Base pair substitutions that replace the targeted allele with an alternative are likely to occur in one in every 10^8^–10^9^ cells, given observed mutation rates per cell division^43^. Additional alterations affecting nearby nucleic or amino acids could interfere with genetic (e.g., CRISPR and RNAi) or protein-targeting approaches. It is also possible that alternative pathways exist for some GEMINI genes, whereby alterations of other genes compensate for inhibition of a GEMINI gene. However, our list of GEMINI genes is highly enriched for components of universal cellular processes, such as DNA, RNA, and protein biogenesis, for which no alternative pathways exist to compensate their loss^44^.

Biomarkers for detection of patients who may benefit from a GEMINI approach are relatively straightforward: one would select patients who are heterozygous for the targeted allele, and for whom the tumor is found to have lost the alternative allele. One consideration is tumor heterogeneity; if the LOH event is present in only part of the tumor, resistance would be expected to arise quickly. However, in prior analyses^43,45–47^, a majority of somatic copy-number alterations, including LOH events, appeared to be clonal, although the fraction of clonal events can be lower in some loci in some tissues^48^. One approach to minimize clonal variation in LOH is to prioritize GEMINI genes that lie on chromosomes or chromosome arms that are characteristically lost early in oncogenesis (e.g., 3p in renal clear cell carcinoma)^37^.

While we show that cells heterozygous for the targeted SNPs of *PRIM1* show no substantial proliferation defects upon ablation of the targeted allele (Figure 2E, 2F), systemic knockout of one allele of an essential gene across all cells in a patient is not likely to be a tractable therapeutic strategy. Thus, potential allele-specific gene editing approaches to leverage GEMINI vulnerabilities in the clinic would rely on a highly cancer cell–specific delivery system to avoid knockout of the targeted allele in normal tissue. While much work remains to achieve the necessary targeting specificity, advances in nanoparticle delivery systems present the possibility of targeting Cas9 DNA, mRNA, or protein in a tumor-specific manner^49–52^. Additionally, *S. pyogenes* Cas9 enzymes with altered^53^ or expanded^54,55^ PAM specificities or CRISPR effectors from other species^53,56–58^ have broadened the total number of targetable loci in the genome and, thereby, the number of targetable variants.

GEMINI vulnerabilities could be also targetable by reversible genetic approaches. Early studies of the GEMINI genes *POLR2A* and *RPA1* achieved allele-specific growth suppression of cancer cells using antisense oligonucleotide^5,59,60^ and RNAi^61^ reagents. Peptide nucleic acids, or PNAs, which can suppress both transcription and translation of target genes, represent another potential allele-specific genetic approach^62–65^. Finally, the RNA-targeting CRISPR effector Cas13 has shown the ability to knock down target genes^66^ and decrease proliferation of cancer cells^67^ in an allele-specific manner. Unlike Cas9, the Cas13 enzyme from *L. wadei* previously used for allele-specific RNA cleavage does not require a downstream PAM-like motif^66^, potentially expanding the number of targetable sites beyond those tractable with DNA-targeting CRISPR effectors. Like Cas9-based modalities, however, Cas13 and other reversible genetic approaches to exploit GEMINI vulnerabilities would require the development of novel delivery systems.

Allele-specific small molecule inhibitors present another attractive possibility for drugging GEMINI vulnerabilities. Allele-specific therapeutics in clinical use include rationally designed drugs (e.g., mutant EGFR inhibitors^68^) as well as those whose genotype-specific effects were identified by pharmacogenomic studies (e.g., warfarin and VKORC1^69^). However, GEMINI vulnerabilities present an additional challenge for allele-specific inhibitor development because most variants in cell-essential genes do not reside in or near an active site (Figure 4C) or other functionally critical protein region. This challenge may be addressed through alternative small-molecule approaches, such as proteolysis-targeting chimera (PROTAC)-mediated degradation^70^. SNPs for which one allele is a cysteine could be prioritized for this approach because of the possibility of engineering a covalent inhibitor^71^.

While we have rigorously validated *PRIM1* and *EXOSC8* as genetic dependencies in cancer, further work is necessary to explore potential therapeutic modalities for targeting them. As an enzyme, PRIM1 represents a potential target for small-molecule drug development. However, while several potential inhibitors of human DNA primase have been proposed, none have yet advanced to clinical stages of development^72^. Efforts to discover additional putative primase inhibitors may benefit from approaches used to target bacterial and viral primases. For instance, a combined fragment-based/virtual screening approach has been used to identify novel inhibitors of the T7 bacteriophage primase. Such an approach may aid PRIM1-targeting lead-compound discovery by eliminating the challenging task of initial primase functional screening for large numbers of compounds^73^.

The residue coded for by *EXOSC8*^rs117135638^ lies on the interface of *EXOSC8* gene product Rrp43 and exosome complex member Mtr3, raising the possibility of developing an allele-specific inhibitor of exosome complex formation. Pharmacologic inhibition of protein-protein binding has generally proven challenging due to the large, often flat, surfaces involved in protein complex formation^74^. However, disruption of several clinically relevant protein-protein interactions has been achieved previously, as illustrated by small-molecule or peptidomimetic inhibitors of p53-Mdm2 binding^75–77^, Notch complex assembly^78^, and herpesvirus DNA polymerase complex assembly^79^. As with *PRIM1*, any such approach to targeting LOH of *EXOSC8* must also overcome the substantial hurdle of achieving allele specificity.

The design of a tractable therapeutic that targets any single GEMINI gene in an allele-specific manner is a substantial challenge. However, the sheer number of potential candidates suggests that some of these GEMINI vulnerabilities may represent viable targets.

### Methods Variant lists

A list of 228,440 potentially targetable variants was downloaded from the Exome Aggregation Consortium (ExAC) database (exac.broadinstitute.org)^9^. Potentially targetable variants were defined as those in the following classes: 3_prime_UTR_variant, 5_prime_UTR_variant, frameshift_variant, inframe_deletion, inframe_insertion, initiator_codon_variant, missense_variant, splice_acceptor_variant, splice_donor_variant, splice_region_variant, stop_gained, stop_lost, stop_retained_variant, synonymous_variant. These variants were filtered to include only PASSing, common variants (global minor allele frequency ≥ 0.01) in genes for which copy number calls were available through the NCI Genomic Data Commons (see below for further details of copy number analyses).

All variant classes were included in the analysis of potential target SNPs for reversible genetic therapeutic approaches. All variant classes except 3_prime_UTR_variant and 5_prime_UTR_variant were included in the determination of variants generating or disrupting an *S. pyogenes* PAM site.

### Genomic analyses of copy number and loss of heterozygosity from TCGA

Patient-derived genome-wide copy number and LOH data were downloaded from the TCGA Pan-Can project via the NCI Genomic Data Commons (https://gdc.cancer.gov/about-data/publications/pancan-aneuploidy) first reported in ^80^. For copy number, gene-level log2 relative data were calculated by GISTIC 2.0, referenced in the output file “all_data_by_genes_whitelisted.tsv”. Copy-loss was defined as log2 relative values ≤-0.1 and copy-neutral was defined as ≥ −0.1.

For LOH calls, ABSOLUTE^81^ was used to generate LOH calls from genomic segments and converted gene-level data from the output file “TCGA_mastercalls.abs_segtabs.fixed.txt”. All subsequent analyses, including genomic frequencies of LOH, were calculated as a per-gene rate.

### Essential gene list

Candidate essential genes were nominated using data from three genome-scale loss of function screens of haploid human cell lines (KBM7 with CRISPR-Cas9 gene inactivation or mutagenized with gene trapping^1^, and pluripotent stem cells with CRISPR-Cas9 gene inactivation^2^). Briefly, all genes that passed a threshold of <10% FDR for a given cell line were included as a candidate essential gene. FDR corrected p-values from the original publications were used for both CRISPR screens; FDR q-values for the KBM7 gene trap scores were calculated using a binomial model (representing equal probability of gene trap inserting in a sense versus anti-sense orientation) and correction for multiple hypotheses using Benjamini and Hochberg. This initial candidate list contained 3431 genes, with 633 scoring as essential in all three screens.

These candidate essential genes were then filtered using CCLE gene copy-number and RNA expression data to determine if loss of function genetic alterations were observed in human cell lines. Genes that met any of the following criteria were excluded: homozygously deleted in >2 cell lines (log2 copy-number < −5); very low RNA expression (< 0.5 RPKM) in >5 cell lines; or homozygously deleted in 1 cell line that also has low RNA expression (<1.0 RPKM). This analysis reduced the list to 2566 candidate essential genes. Genes were then filtered based on mean CERES score from CRISPR knockout screens of 517 cell lines (https://depmap.org/portal/download; derived from the file “gene_effects.csv”)^26^. Genes with CERES scores < −0.4 were excluded, yielding a list of 1499 essential genes. To account for instances in which CCLE copy-number and/or RNA expression data were not available for a particular gene, genes were rescued from the CCLE filter if they scored as essential in two of the three haploid cell line screens and had mean a CERES score < −0.4. This rescue yielded 17 genes, bringing the total number of candidate essential genes to 1516. This list was further filtered to remove genes classified as Tier 1 tumor suppressor genes in the COSMIC Cancer Gene Census (https://cancer.sanger.ac.uk/census/)^3^, yielding a final list of 1482 essential genes. (One essential gene in this list, AK6, was not characterized in TCGA copy-number and LOH data and so was excluded from further analyses.)

### Cell line identification and cell culture

Human cancer cell lines of the appropriate genotypes for *PRIM1* and *EXOSC8* were identified using whole exome sequencing and absolute gene copy number data from the Cancer Cell Line Encyclopedia (https://portals.broadinstitute.org/ccle)^83^. All lines were genotyped for the SNP of interest using Sanger sequencing. Cell lines were maintained in RPMI-1640 supplemented with 10% fetal bovine serum and 1% penicillin. Lines were not assessed for contamination with mycoplasma. No commonly misidentified cell lines defined by the International Cell Line Authentication Committee have been used in these studies.

### Plasmids

lentiCas9-Blast (Addgene plasmid # 52962) and lentiGuide-Puro (Addgene plasmid # 52963) were gifts from Feng Zhang^84^. A Cas9 construct co-expressing GFP and two sgRNAs was a gift from Peter Choi^26^. pLKO.1–TRC cloning vector was a gift from David Root (Addgene plasmid # 10878)^85^.

### CRISPR sgRNAs

To identify target sites for CRISPR-Cas9-mediated knockout, the genetic sequences of *PRIM1* and *EXOSC8* were obtained from the UCSC genome browser (http://genome.ucsc.edu) using the human assembly GRC38/Hg38 (December 2013). The 20 nucleotides upstream of the polymporphic PAM site containing the SNP for each gene constitutes the AS sgRNA for that gene. All other sgRNAs were designed using the CRISPR sgRNA design tool from the Zhang lab (http://crispr.mit.edu). sgRNAs were cloned into the appropriate vector as described previously^84,86^. The sgRNA sequences were as follows:

LacZ: GTTCGCATTATCCGAACCAT
PRIM1 AS: CAGCTCGGGCAGCTCGGTGG
PRIM1 NA: CGCTGGCTCAACTACGGTGG
EXOSC8 AS: CGGAATCTCGATGAACACAG
EXOSC8 NA: ACCGGAATCTCGATGAACAC

### Cell growth assays

Cells were plated in opaque 96-well plates at 500 or 1000 cells per well (Corning) on the indicated day post-lentiviral infection. Cell number was inferred by ATP-dependent luminescence using CellTiter-Glo (Promega) reagent and normalized to the relative luminescence on the day of plating.

### Generation of *PRIM1*-loss and *EXOSC8*-loss cells by CRISPR-Cas9

A Cas9 construct co-expressing GFP and two sgRNAs with target sites flanking *PRIM1* was used to delete a 20.6kb region encoding PRIM1. Cell lines heterozygous for *PRIM1*^rs2277339^ (SNU-C4, SNU-175, and TYK-nu) were transfected with this construct using LipoD293 transfection reagent (SignaGen), and single GFP+ cells were sorted by FACS and plated at low density for single cell cloning or single-cell sorted into 96-well tissue culture plates containing a 50:50 mix of conditioned and fresh RPMI-1640 media, 20% serum, 1% penicillin-streptomycin, and 10µm ROCK inhibitor Y-27632. Clones were expanded and validated by PCR to harbor the 20.6kb deletion encoding *PRIM1*, and the retained allele was genotyped by Sanger sequencing. These clones were designated SNU-175^PRIM1 S/–^, SNU-175^PRIM1 R/–^, SNU-C4^PRIM1 S/–^ for subsequent experiments. Other clones were determined by PCR and Sanger sequencing to retain both *PRIM1* alleles and not to harbor this deletion and were designated as control cell lines for subsequent experiments (SNU-175^PRIM1 R/S^ and SNU-C4^PRIM1 R/S^). The same procedure was employed using a cell line diploid for the EXOSC8^R^ SNP (SW-579) to generate EXOSC8^R/–^ cell lines harboring a 7.1kb deletion and EXOSC8^R/R^ control lines.

The sgRNA sequences were as follows:

PRIM1 upstream: GCGCGGAACTCGCCACGGTA
PRIM1 downstream: CAGAGCTCCTCAAACCATTG
EXOSC8 upstream: GGTTTCTCGGCCGAGCGCCG
EXOSC8 downstream: TGTACCCATCTACTTAAGTT

Primers used to verify gene deletion by PCR were as follows:

PRIM1 deletion genotyping F: ACTGTATGCACCACCACACC
PRIM1 deletion genotyping R: AGTTCACGTGGAGCATCCTT
EXOSC8 deletion genotyping F: TTTGGGGCATACTCATGCTT
EXOSC8 deletion genotyping R: TCCACCTCCAATTATTTGTTCC

### Generation of *EXOSC8* isogenic cell lines by CRISPR-Cas9–mediated HDR editing

Cas9 RNPs and a ssODN repair template were used to edit the EXOSC8^S^ SNP to the EXOSC8^R^ SNP. S. pyogenes Cas9-NLS (Synthego) and an sgRNA (sequence: ACCGGAATCTCGATGAACAC) targeting the *EXOSC8* SNP region (Synthego) were complexed as described previously^87^. DV-90 cells (EXOSC8^S/S^) were nucleofected with resulting RNPs, a 50:50 mix of EXOSC8^S^ and EXOSC8^R^ ssODN (IDT), and a GFP-expressing plasmid (pMAX-GFP) (Lonza). ssODN repair templates contained a synonymous mutation introducing a novel *Mnl1* restriction site for downstream genotyping as well as a silent blocking mutation to prevent repeated Cas9 cleavage. Single GFP+ cells were single-cell sorted by FACS into 96-well tissue culture plates containing a 50:50 mix of conditioned and fresh RPMI-1640 media, 20% serum, 1% penicillin-streptomycin, and 10µm ROCK inhibitor Y-27632. Clones were expanded and evaluated for HDR-mediated editing by PCR and restriction digest, and positive clones were genotyped by next-generation sequencing (MGH DNA Core).

### CRISPR variant sequencing

Cellular pellets were collected from Cas9-stable cells 4 or 18 days post-infection with lentiGuide-Puro virus encoding the indicated sgRNA. Genomic DNA was isolated using a DNAMini kit (Qiagen), and the target region for each gene was amplified by PCR (EMD Millipore). Amplicons were submitted to NGS CRISPR sequencing by the MGH DNA Core. Frameshift and non-inactivating alleles (nonaltered or in-frame indels) were determined manually using the CRISPR variant output file. PCR primer sequences were as follows:

PRIM1 MGH F: GCACAGAAGGCGCTTCATA
PRIM1 MGH R: CGCCAATTCCTGTGGTAATC
EXOSC8 F: AGCTGCAGAGTGTTTCTTTCA
EXOSC8 R: AGAGCAAAGTAAATGAAAAGCCCAA

### Western blotting

Cells were washed in ice-cold PBS and lysed in 1x RIPA buffer (10 mM Tris-Cl Ph 8.0, 1 mM EDTA, 1% Triton X-100, 0.1% SDS and 140 mM NaCl) supplemented with 1x protease and phosphatase inhibitor cocktail (PI-290, Boston Bioproducts). Lysates were sonicated in a bioruptor (Diagenode) for 5 min (medium intensity) and cleared by centrifugation at 15,000 x g for 15 min at 4°C. Proteins were electrophoresed on polyacrylamide gradient gels (Life Technologies) and detected by chemiluminescence (Bio-rad). Antibodies used were as follows:

EXOSC8: Proteintech #11979-1-AP
PRIM1: Cell Signaling Technology #4725
Vinculin: Sigma #V9131

### shRNA sequences

pLKO.1 GFP shRNA (target sequence: GCAAGCTGACCCTGAAGTTCAT) was a gift from David Sabatini (Addgene plasmid # 30323)^88^. Lentiviral expression constructs for non-allele specific shRNA-mediated suppression of *PRIM1* were obtained through the Broad Institute of MIT and Harvard Genomic Perturbation Platform (https://portals.broadinstitute.org/gpp/public/). The names, clone IDs, and target sequences used in our studies are as follows:

shPRIM1 (TRCN0000275194): AGCATCGTCTCTGGGTATATT
TRCN0000151860: CCGAGCTGCTTAAACTTTATT
TRCN0000275194: AGCATCGTCTCTGGGTATATT
TRCN0000275195: GATTGATATAGGCGCAGTATA
TRCN0000275196: CCGAGCTGCTTAAACTTTATT

The following allele-specific shRNA sequences were cloned into the vector pLKO.1 as described previously^85^:

*PRIM1*^rs2277339^ major-allele (T) targeting:

sh3T: TCAATGGAGACGTTTGACC
sh4T: CAATGGAGACGTTTGACCC
sh5T: AATGGAGACGTTTGACCCC
sh6T: ATGGAGACGTTTGACCCCA
sh7T: TGGAGACGTTTGACCCCAC
sh8T: GGAGACGTTTGACCCCACC
sh9T: GAGACGTTTGACCCCACCG
sh10T: AGACGTTTGACCCCACCGA
sh11T: GACGTTTGACCCCACCGAG
sh16T: TTGACCCCACCGAGCTGCC

*PRIM1*^rs2277339^ minor-allele (G) targeting:

sh3G: TCAATGGAGACGTTTGCCC
sh4G: CAATGGAGACGTTTGCCCC
sh5G: AATGGAGACGTTTGCCCCC
sh6G: ATGGAGACGTTTGCCCCCA
sh7G: TGGAGACGTTTGCCCCCAC
sh8G: GGAGACGTTTGCCCCCACC
sh9G: GAGACGTTTGCCCCCACCG
sh10G: AGACGTTTGCCCCCACCGA
sh11G: GACGTTTGCCCCCACCGAG
sh16G: TTGCCCCCACCGAGCTGCC

### Quantitative and reverse transcription PCR

RNA was extracted using the RNeasy extraction kit (Qiagen) and subjected to on-column DNase treatment. cDNA was synthesized with the Superscript II Reverse Transcriptase kit (Life Technologies) with no reverse transcriptase samples serving as negative controls. Gene expression was quantified by Power Sybr Green Master Mix (Applied Biosystems). *PRIM1* expression values were normalized to vinculin (*VCL*) and the fold change calculated by the DDCt method. Primers used in our studies are as follows:

PRIM1-F: GCTCAACTACGGTGGAGTGAT
PRIM1-R: GGTTGTTGAAGGATTGGTAGCG
VCL-F: CGCTGAGGTGGGTATAGGTG
VCL-R: TTGGATGGCATTAACAGCAG.

### Calculations of theoretical patient numbers

To determine number of patients that could benefit from a therapeutic approach targeting each GEMINI vulnerability, we used the following formula:

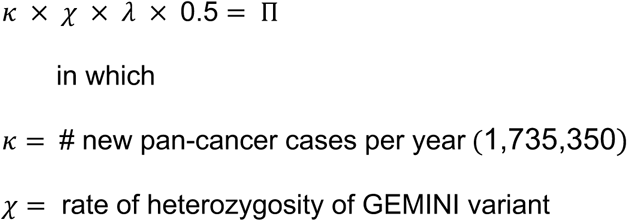

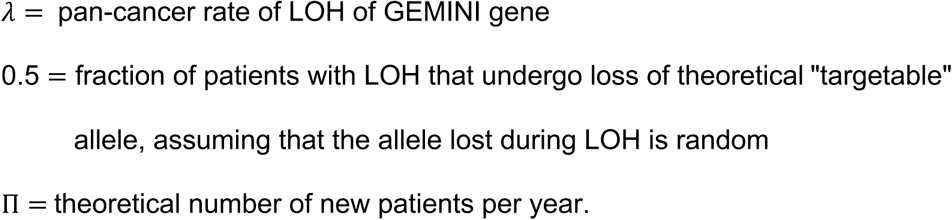

Estimate of new pan-cancer cases per year derived from SEER Cancer Statistics Review^89^. Rate of heterozygosity estimated using 2pq from Hardy-Weinberg equation^90,91^.

### DEMETER2 analyses

For a detailed description of the screening and analysis methodology used to generate DEMETER2 scores, please see ^28^. Briefly, DEMETER2 generates an absolute dependency score for each gene suppressed in each cell line. A score of 0 signifies no dependency and a score of 1 signifies a strong dependency as estimated by scaling the effect to a panel of known pan-essential genes.

DEMETER2 scores were obtained from the Cancer Depencency Map Portal (https://depmap.org/portal/download/) using the file “D2_combined_gene_dep_scores.csv”. We classified genes that exhibited a median DEMETER2 score of ≤ −0.5 across all cell lines as moderately strong dependencies.

### canSAR protein annotation

The canSAR protein annotation tool (cansar.icr.ac.uk) was run on a list of 741 unique genes containing 1749 insertion, deletion, and missense variants.

Structures with >90% sequence homology were included in structural druggability and chemical matter analyses.

### Determination of structures corresponding to variants

To determine which variants were present in PDB, DNA sequences (30mer) encapsulating 1749 insertion, deletion, and missense variants were translated in all 6 frames using the Bio.Seq Python module. Output was blasted using the Bio.Blast Python module against the PDB database with E-value thresholds of 0.001 or less, resulting in hits for 267 variants. We manually curated these structures to verify the presence of the variant within the PDB file and eliminated structures for which correspondence between the PDB protein sequence, ExAC amino acid prediction, and UCSC Genome Browser amino acid sequence was inconclusive. This curation yielded 153 protein-altering variants in proteins with homologous molecular structures.

Visual scoring was performed on 81 protein-altering variants that lie in X-ray crystal structures. Variants were scored using the following scale: 0 = no clear pockets on the protein surface, 1 = SNP far from pocket on protein surface, 2 = SNP near pocket on protein surface, 3 = SNP in pocket on protein surface, 4 = SNP near pocket containing small molecule.

### Data Availability

The datasets analyzed during the current study are available in the following repositories:

Exome Aggregation Consortium, http://exac.broadinstitute.org/downloads NCI Genomic Data Commons: TCGA copy number and LOH data, https://gdc.cancer.gov/about-data/publications/pancan-aneuploidy; CCLE whole exome sequencing data, https://portal.gdc.cancer.gov/legacy-archive/search/f

Cancer Cell Line Encyclopedia Portal: https://portals.broadinstitute.org/ccle

Cancer Depencency Map Portal: https://depmap.org/portal/download/

## Author Contributions

CAN, WJG: designed and performed experiments, analyzed data, and prepared the manuscript; MSB, JPB, HW: designed and performed experiments and analyzed data; LMU, NC: designed and performed experiments; JAK: analyzed data and provided technical support and conceptual advice; AB, GG: analyzed data; ADC: analyzed data and provided technical support and conceptual advice; BRP: designed and performed experiments, analyzed data, prepared the manuscript, and provided technical support and conceptual advice; SDP, RB: analyzed data, prepared the manuscript, and provided technical support and conceptual advice.

Supplementary Figure 1: Genomic landscape of essential genes, LOH, and GEMINI genes.

Supplementary Figure 2: Additional *PRIM1* statistics and growth curve replicates.

Supplementary Figure 3: Genotyping of *PRIM1*^rs2277339^ and *EXOSC8*^rs117135638^ isogenic cell lines.

Supplementary Figure 4: Additional *EXOSC8* statistics and growth curve replicates.

Supplementary Figure 5: Growth effects of allele-specific shRNAs targeting *PRIM1*_rs2277339_.

Supplementary Table 1: Cell-essential genes.

Supplementary Table 2: DAVID Biological Process GO Term enrichment analysis for cell-essential genes.

Supplementary Table 3: GEMINI vulnerabilities.

Supplementary Table 4: DAVID Biological Process GO Term enrichment analysis for GEMINI genes.

Supplementary Table 5: Annotation of protein-altering GEMINI vulnerabilities.

## References

1. Boehm, J. S., Hession, M. T., Bulmer, S. E. & Hahn, W. C. Transformation of Human and Murine Fibroblasts without Viral Oncoproteins. Mol. Cell. Biol. 25, 6464–6474 (2005).

2. Vogelstein, B. et al. Cancer Genome Landscapes. Science (80-. ). 339, 1546–1558 (2013).

3. Martincorena, I. et al. Universal Patterns of Selection in Cancer and Somatic Tissues. Cell 171, 1029–1041 (2017).

4. Fluiter, K., Housman, D., Ten Asbroek, a L. M. a & Baas, F. Killing cancer by targeting genes that cancer cells have lost: Allele-specific inhibition, a novel approach to the treatment of genetic disorders. Cell. Mol. Life Sci. 60, 834–843 (2003).

5. Basilion, J. P. et al. Selective Killing of Cancer Cells Based on Loss of Heterozygosity and Normal Variation in the Human Genome: A New Paradigm for Anticancer Drug Therapy. Mol. Pharmacol. 56, 359–369 (1999).

6. ten Asbroek, A. L. M. A., Fluiter, K., van Groenigen, M., Nooij, M. & Baas, F. Polymorphisms in the large subunit of human RNA polymerase II as target for allele-specific inhibition. Nucleic Acids Res. 28, 1133–1138 (2000).

7. Huang, D. W., Sherman, B. T. & Lempicki, R. A. Bioinformatics enrichment tools: Paths toward the comprehensive functional analysis of large gene lists. Nucleic Acids Res. 37, 1–13 (2009).

8. Huang, D. W., Sherman, B. T. & Lempicki, R. A. Systematic and integrative analysis of large gene lists using DAVID bioinformatics resources. Nat. Protoc. 4, 44–57 (2009).

9. Lek, M. et al. Analysis of protein-coding genetic variation in 60,706 humans. Nature 536, 285–291 (2016).

10. Beroukhim, R. et al. The landscape of somatic copy-number alteration across human cancers. Nature 463, 899–905 (2010).

11. Zack, T. I. et al. Pan-cancer patterns of somatic copy number alteration. Nat. Genet. 45, 1134–1140 (2013).

12. Holland, A. J. & Cleveland, D. W. Boveri revisited: Chromosomal instability, aneuploidy and tumorigenesis. Nat. Rev. Mol. Cell Biol. 10, 478–487 (2009).

13. Courtney, D. G. et al. CRISPR/Cas9 DNA cleavage at SNP-derived PAM enables both in vitro and in vivo KRT12 mutation-specific targeting. Gene Ther. 23, 108–112 (2016).

14. Shin, J. W. et al. Permanent inactivation of Huntington’s disease mutation by personalized allele-specific CRISPR/Cas9. Hum. Mol. Genet. 25, 4566– 4576 (2016).

15. Christie, K. A. et al. Towards personalised allele-specific CRISPR gene editing to treat autosomal dominant disorders. Sci. Rep. 7, 1–11 (2017).

16. Mojica, F. J. M., Díez-Villaseñor, C., García-Martínez, J. & Almendros, C. Short motif sequences determine the targets of the prokaryotic CRISPR defence system. Microbiology 155, 733–740 (2009).

17. Hsu, P. D. et al. DNA targeting specificity of RNA-guided Cas9 nucleases. Nat. Biotechnol. 31, 827–832 (2013).

18. Lucchini, G., Francesconi, S., Foiani, M., Badaracco, G. & Plevani, P. Yeast DNA polymerase–DNA primase complex: cloning of PRI 1, a single essential gene related to DNA primase activity. EMBO J. 6, 737–742 (1987).

19. Francesconi, S. et al. Mutations in conserved yeast DNA primase domains impair DNA replication in vivo. Proc. Natl. Acad. Sci. U. S. A. 88, 3877– 3881 (1991).

20. Frick, D. N. & Richardson, C. C. DNA Primases. Annu. Rev. Biochem. 70, 39–80 (2001).

21. Mitchell, P., Petfalski, E., Shevchenko, A., Mann, M. & Tollervey, D. The Exosome: A Conserved Eukaryotic RNA Processing Complex Containing Multiple 3’-->5’ Exoribonucleases. Cell 91, 457–466 (1997).

22. Schmid, M. & Jensen, T. H. The exosome: a multipurpose RNA-decay machine. Trends Biochem. Sci. 33, 501–510 (2008).

23. Kilchert, C., Wittmann, S. & Vasiljeva, L. The regulation and functions of the nuclear RNA exosome complex. Nat. Rev. Mol. Cell Biol. 17, 227–239 (2016).

24. Jiang, W., Bikard, D., Cox, D., Zhang, F. & Marraffini, L. a. RNA-guided editing of bacterial genomes using CRISPR-Cas systems. Nat. Biotechnol. 31, 233–9 (2013).

25. Nijhawan, D. et al. Cancer Vulnerabilities Unveiled by Genomic Loss. Cell 150, 842–854 (2012).

26. Paolella, B. R. et al. Copy-number and gene dependency analysis reveals partial copy loss of wild-type SF3B1 as a novel cancer vulnerability. Elife 6, 1–35 (2017).

27. Tsherniak, A. et al. Defining a Cancer Dependency Map. Cell 170, 564– 576 (2017).

28. McFarland, J. M. et al. Improved estimation of cancer dependencies from large-scale RNAi screens using model-based normalization and data integration. bioRxiv 305656 (2018). doi:10.1101/305656

29. Halling-Brown, M. D., Bulusu, K. C., Patel, M., Tym, J. E. & Al-Lazikani, B. canSAR: an integrated cancer public translational research and drug discovery resource. Nucleic Acids Res. 40, 947–956 (2012).

30. Tym, J. E. et al. canSAR: An updated cancer research and drug discovery knowledgebase. Nucleic Acids Res. 44, D938–D943 (2016).

31. Enzyme Nomenclature 1992. (Academic Press, 1992).

32. Berman, H. M. et al. The Protein Data Bank. Nucleic Acids Res. 28, 235– 242 (2000).

33. Nijman, S. M. B. Synthetic lethality: General principles, utility and detection using genetic screens in human cells. FEBS Lett. 585, 1–6 (2011).

34. Hartwell, L. H., Szankasi, P., Roberts, C. J., Murray, A. W. & Stephen H. Friend. Integrating Genetic Approaches into the Discovery of Anticancer Drugs. Science (80-. ). 278, 1064–1068 (1997).

35. McLornan, D. P., List, A. & Mufti, G. J. Applying Synthetic Lethality for the Selective Targeting of Cancer. N. Engl. J. Med. 371, 1725–1735 (2014).

36. Muller, F. L., Aquilanti, E. A. & DePinho, R. A. Collateral Lethality: A New Therapeutic Strategy in Oncology. Trends in Cancer 1, 161–173 (2015).

37. Muller, F. L. et al. Passenger Deletions Generate Therapeutic Vulnerabilities in Cancer. Nature 488, 337–342 (2013).

38. Helming, K. C. et al. ARID1B is a specific vulnerability in ARID1A-mutant cancers. Nat. Med. 20, 251–254 (2014).

39. Dey, P. et al. Genomic deletion of malic enzyme 2 confers collateral lethality in pancreatic cancer. Nature 542, 119–123 (2017).

40. Viswanathan, S. R. et al. Genome-scale analysis identifies paralog lethality as a vulnerability of chromosome 1p loss in cancer. Nat. Genet. 50, 937– 9431 (2018).

41. Aksoy, B. A. et al. Prediction of individualized therapeutic vulnerabilities in cancer from genomic profiles. Bioinformatics 30, 2051–2059 (2014).

42. Bailey, M. H. et al. Comprehensive Characterization of Cancer Driver Genes and Mutations. Cell 173, 371–385 (2018).

43. Wang, Y. et al. Clonal evolution in breast cancer revealed by single nucleus genome sequencing. Nature 512, 155–160 (2014).

44. Liu, G. et al. Gene Essentiality Is a Quantitative Property Linked to Cellular Evolvability. Cell 163, 1388–1399 (2015).

45. Kim, T. et al. Subclonal Genomic Architectures of Primary and Metastatic Colorectal Cancer Based on Intratumoral Genetic Heterogeneity. Clin. Cancer Res. 21, 4461–4473 (2015).

46. Gibson, W. J. et al. The genomic landscape and evolution of endometrial carcinoma progression and abdominopelvic metastasis. Nat. Genet. 48, 848–855 (2016).

47. Jamal-Hanjani, M. et al. Tracking the Evolution of Non–Small-Cell Lung Cancer. N. Engl. J. Med. 376, 2109–2121 (2017).

48. McGranahan, N. et al. Allele-Specific HLA Loss and Immune Escape in Lung Cancer Evolution. Cell 171, 1259–1271.e11 (2017).

49. Wang, Y. et al. Systemic Delivery of Modified mRNA Encoding Herpes Simplex Virus 1 Thymidine Kinase for Targeted Cancer Gene Therapy. Mol. Ther. 21, 358–367 (2013).

50. Wang, M., Alberti, K., Sun, S., Arellano, C. L. & Xu, Q. Combinatorially Designed Lipid-like Nanoparticles for Intracellular Delivery of Cytotoxic Protein for Cancer Therapy. Angew. Chemie - Int. Ed. 53, 2893–2898 (2014).

51. Sun, W. et al. Self-Assembled DNA Nanoclews for the Efficient Delivery of CRISPR-Cas9 for Genome Editing. Angew. Chemie - Int. Ed. 54, 12029– 12033 (2015).

52. Wang, H.-X. et al. Nonviral gene editing via CRISPR/Cas9 delivery by membrane-disruptive and endosomolytic helical polypeptide. Proc. Natl. Acad. Sci. 115, 4903–4908 (2018).

53. Kleinstiver, B. P. et al. Engineered CRISPR-Cas9 nucleases with altered PAM specificities. Nature (2015). doi:10.1038/nature14592

54. Hu, J. H. et al. Evolved Cas9 variants with broad PAM compatibility and high DNA specificity. Nature 556, 57–63 (2018).

55. Nishimasu, H. et al. Engineered CRISPR-Cas9 nuclease with expanded targeting space. Science (80-. ). 9129, eaas9129 (2018).

56. Hou, Z. et al. Efficient genome engineering in human pluripotent stem cells using Cas9 from Neisseria meningitidis. PNAS 110, 15644–15649 (2013).

57. Kim, E. et al. In vivo genome editing with a small Cas9 orthologue derived from Campylobacter jejuni. Nat. Commun. 8, 1–12 (2017).

58. Zetsche, B. et al. Cpf1 Is a Single RNA-Guided Endonuclease of a Class 2 CRISPR-Cas System. Cell 163, 759–771 (2015).

59. Fluiter, K. et al. Tumor Genotype-specific Growth Inhibition in Vivo by Antisense Oligonucleotides against a Polymorphic Site of the Large Subunit of Human RNA Polymerase II. Cancer Res. 62, 2024–2028 (2002).

60. Fluiter, K. et al. In vivo tumor growth inhibition and biodistribution studies of locked nucleic acid (LNA) antisense oligonucleotides. Nuc 31, 953–962 (2003).

61. Mook, O. R. F., Baas, F., de Wissel, M. B. & Fluiter, K. Allele-specific cancer cell killing in vitro and in vivo targeting a single-nucleotide polymorphism in POLR2A. Cancer Gene Ther. 16, 532–538 (2009).

62. Hanvey, J. C. et al. Antisense and antigene properties of peptide nucleic acids. Science (80-. ). 258, 1481–1485 (1992).

63. Nielsen, P. E., Egholm, M. & Buchardt, O. Sequence-specific transcription arrest by peptide nucleic acid bound to the DNA template strand. Gene 149, 139–145 (1994).

64. Gambacorti-Passerini, B. C. et al. In Vitro Transcription and Translation Inhibition by Anti-Promyelocytic Leukemia (PML)/Retinoic Acid Receptor Alpha and Anti-PML Peptide Nucleic Acid. Blood 88, 1411–1417 (1996).

65. Egholm, M. et al. PNA hybridizes to complementary oligonucleotides obeying the Watson–Crick hydrogen-bonding rules. Nature 365, 566–568 (1993).

66. Abudayyeh, O. O. et al. RNA targeting with CRISPR–Cas13. Nature 550, 280–284 (2017).

67. Zhao, X. et al. A CRISPR-Cas13a system for efficient and specific therapeutic targeting of mutant KRAS for pancreatic cancer treatment. Cancer Lett. 431, 171–181 (2018).

68. Sullivan, I. & Planchard, D. Next-Generation EGFR Tyrosine Kinase Inhibitors for Treating EGFR-Mutant Lung Cancer beyond First Line. Front. Med. 3, 1–13 (2017).

69. Rost, S. et al. Mutations in VKORC1 cause warfarin resistance and multiple coagulation factor deficiency type 2. Nature 427, 537–541 (2004).

70. Toure, M. & Crews, C. M. Small-molecule PROTACS: New approaches to protein degradation. Angew. Chemie - Int. Ed. 55, 1966–1973 (2016).

71. Lonsdale, R. & Ward, R. A. Structure-based design of targeted covalent inhibitors. Chem. Soc. Rev. 47, 3816–3830 (2018).

72. Boudet, J., Devillier, J. C., Allain, F. H. T. & Lipps, G. Structures to complement the archaeo-eukaryotic primases catalytic cycle description: What’s next? Comput. Struct. Biotechnol. J. 13, 339–351 (2015).

73. Ilic, S. et al. Identification of DNA primase inhibitors via a combined fragment-based and virtual screening. Sci. Rep. 6, 1–10 (2016).

74. Bakail, M. & Ochsenbein, F. Targeting protein-protein interactions, a wide open field for drug design. Comptes Rendus Chim. 19, 19–27 (2016).

75. Lao, B. B. et al. Rational design of topographical helix mimics as potent inhibitors of protein-protein interactions. J. Am. Chem. Soc. 136, 7877– 7888 (2014).

76. Vassilev, L. T. et al. In Vivo Activation of the p53 Pathway by Small-MoleculeAntagonists of MDM2. Science (80-. ). 303, 844–848 (2004).

77. Fasan, R. et al. Using a β-hairpin to mimic an α-helix: Cyclic peptidomimetic inhibitors of the p53-HDM2 protein-protein interaction. Angew. Chemie - Int. Ed. 43, 2109–2112 (2004).

78. Moellering, R. E. et al. Direct inhibition of the NOTCH transcription factor complex. Nature 462, 182–188 (2009).

79. Loregian, A., Marsden, H. S. & Palu, G. Protein-protein interactions as targets for antiviral chemotherapy. Rev. Med. Virol. 12, 239–262 (2002).

80. Taylor, A. M. et al. Genomic and Functional Approaches to Understanding Cancer Aneuploidy. Cancer Cell 33, 676–689 (2018).

81. Carter, S. L. et al. Absolute quantification of somatic DNA alterations in human cancer. Nat. Biotechnol. 30, 413–421 (2012).

82. Wang, T. et al. Identification and characterization of essential genes in the human genome. 1–10 (2015). doi:10.1126/science.aac7041

83. Barretina, J. et al. The Cancer Cell Line Encyclopedia enables predictive modelling of anticancer drug sensitivity. Nature 483, 603–307 (2012).

84. Sanjana, N. E., Shalem, O. & Zhang, F. Improved vectors and genome-wide libraries for CRISPR screening. Nat Methods 11, 783–784 (2014).

85. Moffat, J. et al. A Lentiviral RNAi Library for Human and Mouse Genes Applied to an Arrayed Viral High-Content Screen. Cell 124, 1283–1298 (2006).

86. Shalem, O. et al. Genome-Scale CRISPR-Cas9 Knockout. Science 343, 84–88 (2014).

87. Richardson, C. D., Ray, G. J., DeWitt, M. A., Curie, G. L. & Corn, J. E. Enhancing homology-directed genome editing by catalytically active and inactive CRISPR-Cas9 using asymmetric donor DNA. Nat. Biotechnol. 34, 339–344 (2016).

88. Sancak, Y. et al. The Rag GTPases bind raptor and mediate amino acid signaling to mTORC1. Science (80-. ). 320, 1496–1501 (2008).

89. Noone, A. et al. SEER Cancer Statistics Review, 1975-2015. National Cancer Institute (2017). Available at: https://seer.cancer.gov/csr/1975_2015/,.

90. Hardy, G. H. Mendelian Proportions in a Mixed Population. Science 28, 49–50 (1908).

91. Weinberg, W. Uber den Nachweis der Vererbung beim Menschen. Jahreshefte des Vereins für vaterländische Naturkd. Württemberg. 64, 368–382 (1908).

92. Vaithiyalingam, S. et al. Insights into eukaryotic primer synthesis from structures of the p48 subunit of human DNA primase. J. Mol. Biol. 426, 558–569 (2014).

93. Liu, Q., Greimann, J. C. & Lima, C. D. Reconstitution, Activities, and Structure of the Eukaryotic RNA Exosome. Cell 127, 1223–1237 (2006).

