## Supplementary Figures for "Loss of heterozygosity of essential genes represents a widespread class of potential cancer vulnerabilities"

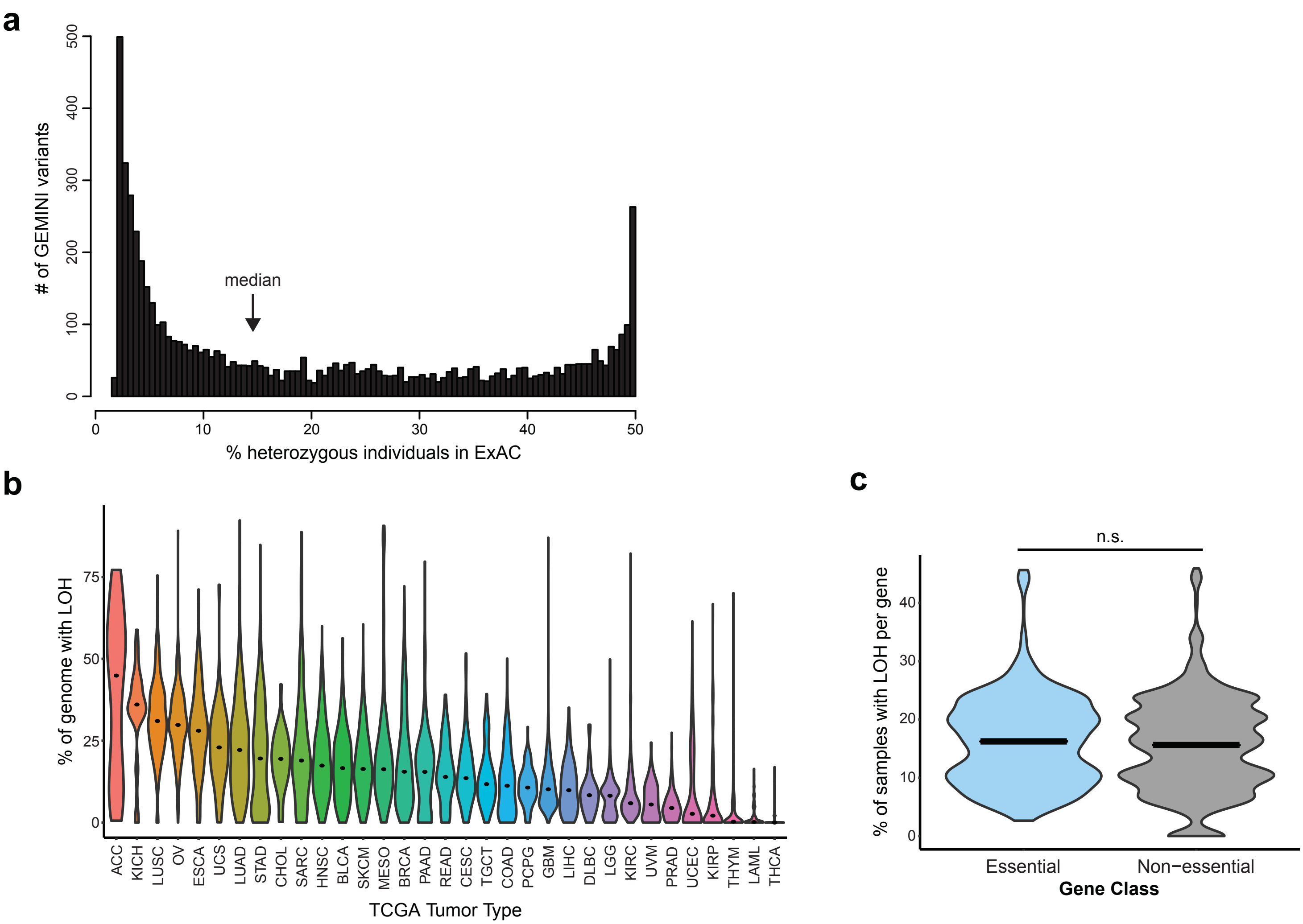


**Supplementary Figure 1.**
**a.** Number of GEMINI variants (vertical axis) plotted against the fraction of individuals that are heterozygous for each variant in the ExAC cohort (horizontal axis). Bin width = 0.5%. **b.** Violin plot of the percent of genome affected by LOH across 33 TCGA tumor types. Tumor types are indicated by TCGA abbreviations (see https://gdc.cancer.gov/resources-tcga-users/tcga-code-tables/tcga-study-abbreviations). Plot width represents relative sample density and dots indicate median values. **c.** Violin plot demonstrating the rate of LOH for essential (blue) and non-essential genes (grey). Essential genes do not have a significantly higher rate of LOH (One-tailed Student's t-test, p=1 [n.s. = not significant]). Intersecting lines indicate median values.

**a**

| *PRIM1*$^{rs2277339}$ | |
|---|---|
| position | chr12:57146069 |
| variant | T/G |
| consequence | missense variant (D5A) |
| minor allele frequency | 0.177 |
| predicted frequency of heterozygosity | 0.291 |
| pan-cancer LOH frequency | 0.089 |
| theoretical patients per year | 22,470 |

**b**

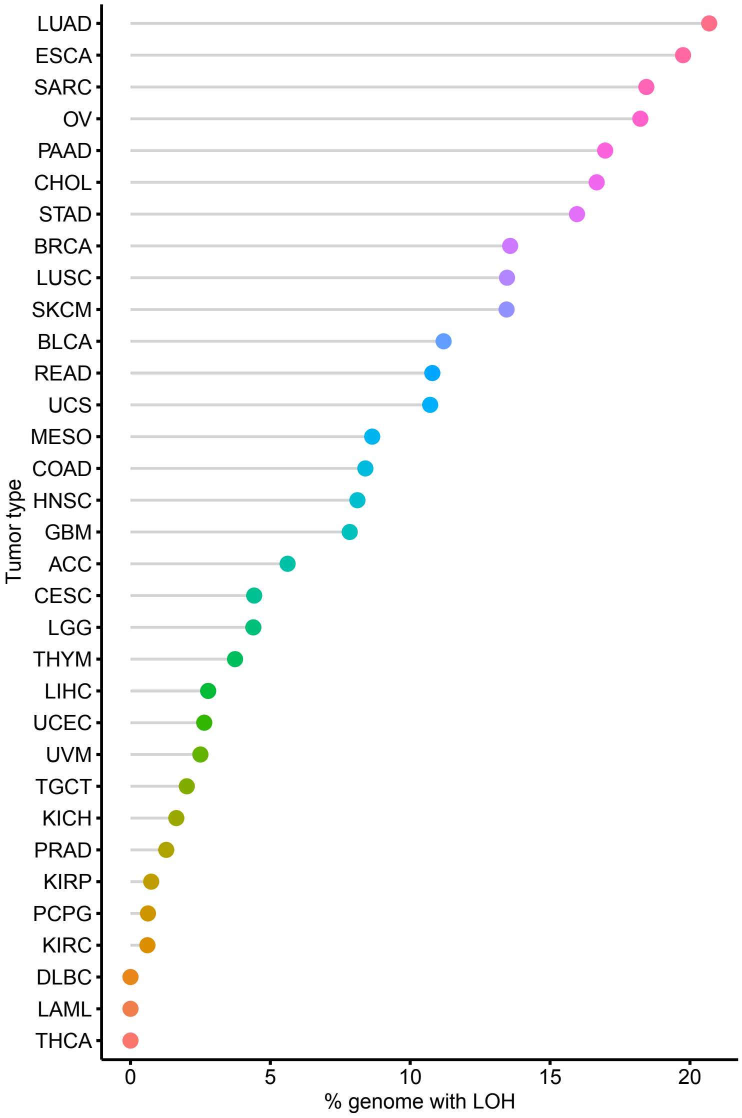


**Supplementary Figure 2.**
**a.** Table of *PRIM1*$^{rs2277339}$ SNP statistics, including calculation of theoretical number of treatable patients per year (see Methods). **b.** Cleveland dot plot of the rate of *PRIM1* LOH across 33 TCGA tumor types. Tumor types are indicated by TCGA abbreviations (see https://gdc.cancer.gov/resources-tcga-users/tcga-code-tables/tcga-study-abbreviations).

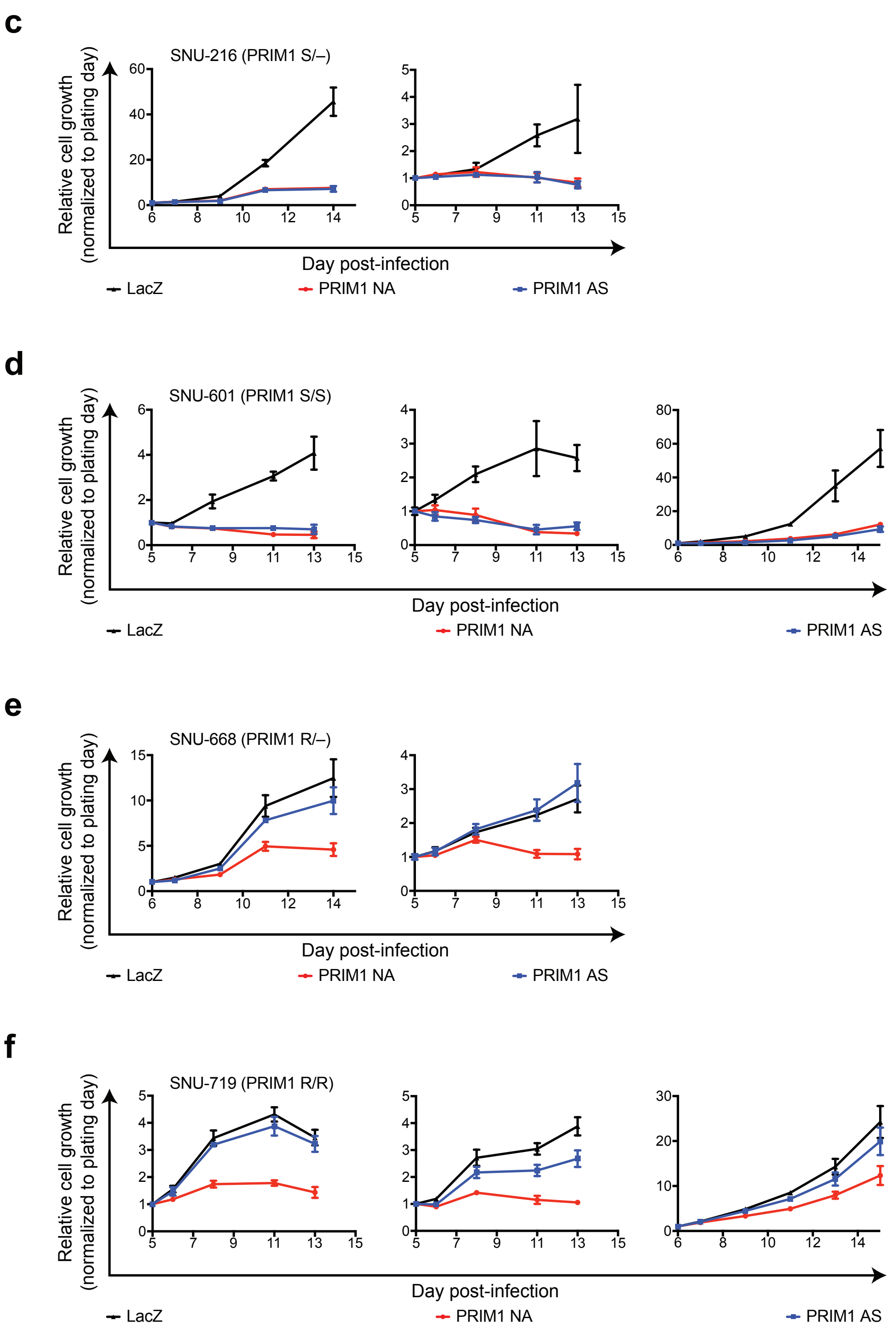


**Supplementary Figure 2.**

**c.–f.** Growth of indicated patient-derived cell lines expressing LacZ (black), PRIM1 NA (red), or PRIM1 AS (blue) sgRNA as measured by CellTiter-Glo luminescence, relative to day of assay plating. n = 5 technical replicates; error bars represent s.d.

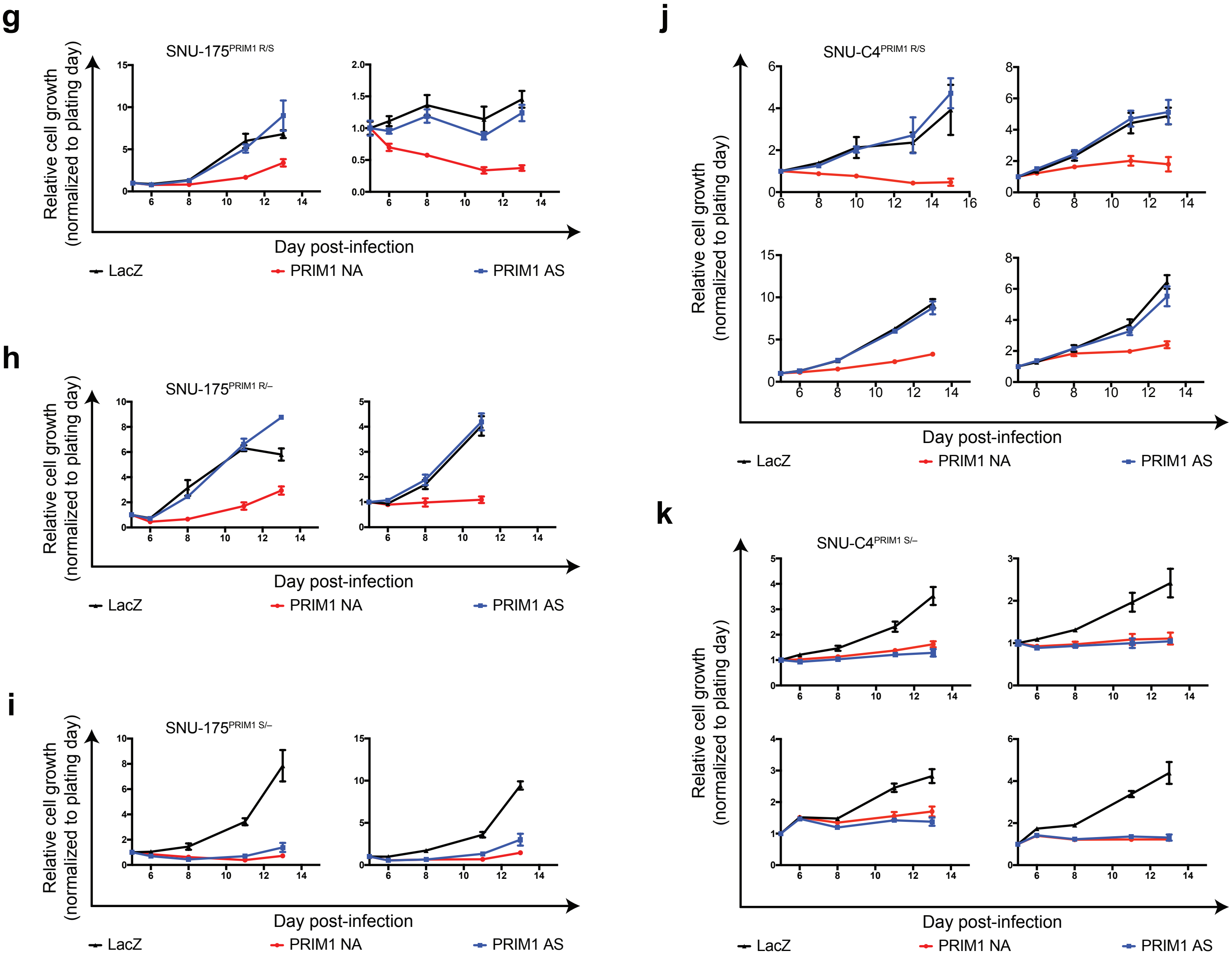


**Supplementary Figure 2.**

**g.–k.** Growth of indicated isogenic cell lines expressing LacZ (black), PRIM1 NA (red), or PRIM1 AS (blue) sgRNA as measured by CellTiter-Glo luminescence, relative to day of assay plating. n = 5 technical replicates; error bars represent s.d.

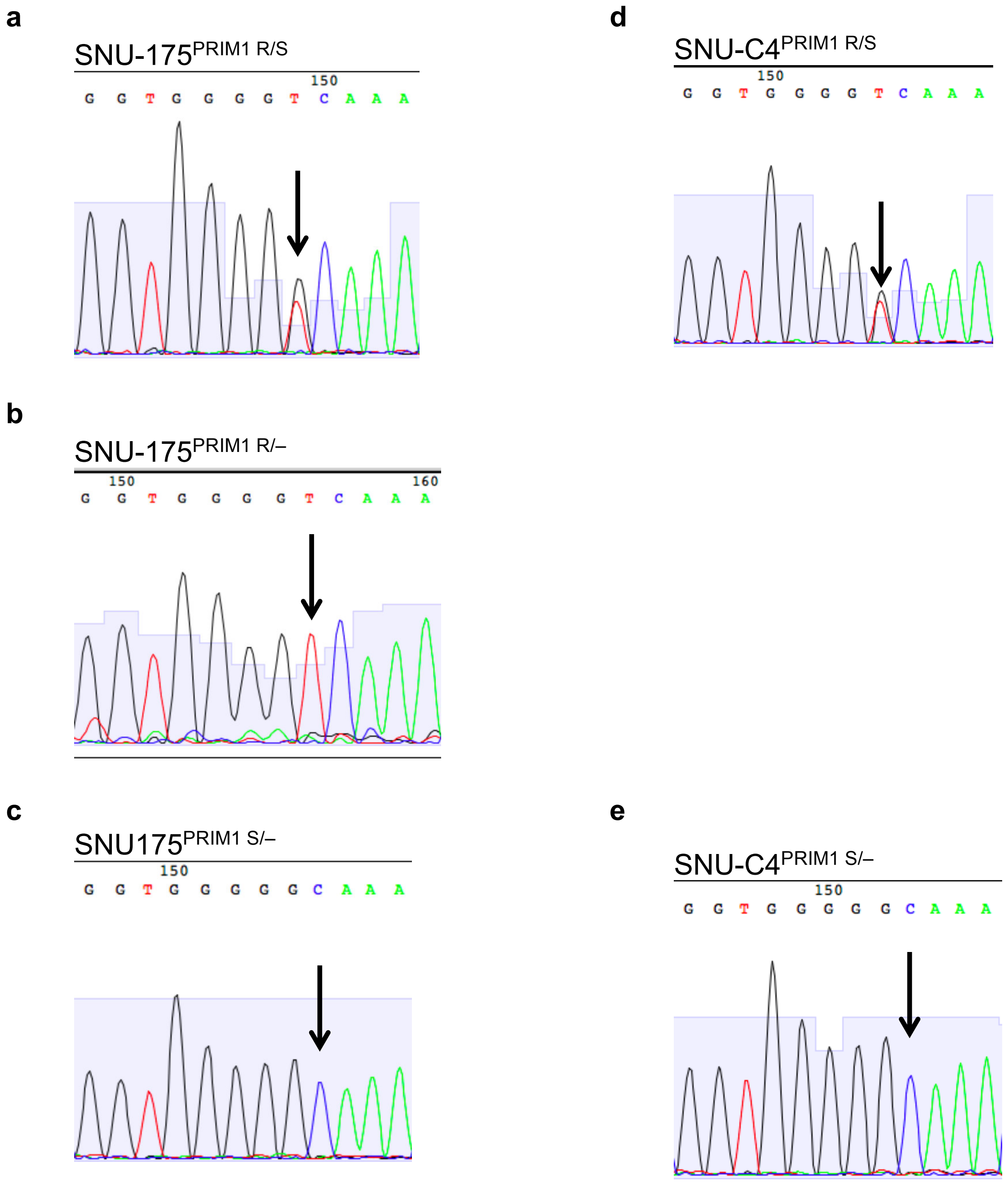


**Supplementary Figure 3.**
**a.–e.** Genotyping of *PRIM1*$^{rs2277339}$ locus in indicated isogenic cell lines by Sanger sequencing. Sequencing peaks representing SNP loci indicated by black arrow.

resistant (R) allele
sensitive (S) allele
editing sgRNA blocking mutation
synonymous mutation introducing MnlI cut site
insertion
deletion

**f**

DV-90$^{EXOSC8\ R/R}$

*Reference*
```
AGCTGCAGAGTGTTTCTTTCAGTTCCTAATGTGGATCTACCACCCCTGTGTTCATCGAGA
```
*Edited R allele, 88,509 read pairs*
```
AGCTGCAGAGTGTTTCTTTCAGTTCCTAATGTGGATCTACCTCATCTGTGTtcatcgaga
***************************************** *  **************
```

**g**

DV-90$^{EXOSC8\ R/\Delta}$

*Reference*
```
AGCTGCAGAGTGTTTCTTTCAGTTCCTAATGTGGATCTACCACCCCTGTG-TTCATCGAG
```
*Edited R allele, 44,454 read pairs*
```
AGCTGCAGAGTGTTTCTTTCAGTTCCTAATGTGGATCTACCTCATCTGTG-Ttcatcgag
```
*Disrupted S allele, 42,169 read pairs*
```
AGCTGCAGAGTGTTTCTTTCAGTTCCTAATGTGGATCTACCACCCCTGTGTTtcatcgag
***************************************** *  ***** *********
```

**h**

DV-90$^{EXOSC8\ S/S}$

*Reference*
```
AGCTGCAGAGTGTTTCTTTCAGTTCCTAATGTGGATCTACCACCCCTGTGTTCATCGAGA
```
*Edited S allele, 54,799 read pairs*
```
AGCTGCAGAGTGTTTCTTTCAGTTCCTAATGTGGATCTACCTCCGCTGTGTtcatcgaga
***************************************** ** ***************
```

**i**

DV-90$^{EXOSC8\ S/\Delta}$

*Reference*
```
AGCTGCAGAGTGTTTCTTTCAGTTCCTAATGTGGATCTACCACCCCTGTGTTCATCGAGA
```
*Disrupted S allele, 22,831 read pairs*
```
AGCTGCAGAGTGTTTCTTTCAGTTCCTAATGTGGATCTACCACCCCTGTG-TCAtcgaga
```
*Edited S allele, 23,409 read pairs*
```
AGCTGCAGAGTGTTTCTTTCAGTTCCTAATGTGGATCTACCTCCGCTGTGTTCAtcgaga
***************************************** ** ***** *********
```

**Supplementary Figure 3.**
**f.–i.** Genotyping of *EXOSC8*$^{rs117135638}$ locus in indicated isogenic cell lines by next-generation sequencing. Relevant sequence features indicated by font or highlighting color. Off-target reads and species representing less than 0.05% of total read pairs removed for clarity.

**a**

| *EXOSC8*rs117135638 | |
|---|---|
| position | 13:37580078 |
| variant | C/A |
| consequence | missense variant (P87H) |
| minor allele frequency | 0.011 |
| predicted frequency of heterozygosity | 0.021 |
| pan-cancer LOH frequency | 0.291 |
| theoretical patients per year | 5307 |

**b**

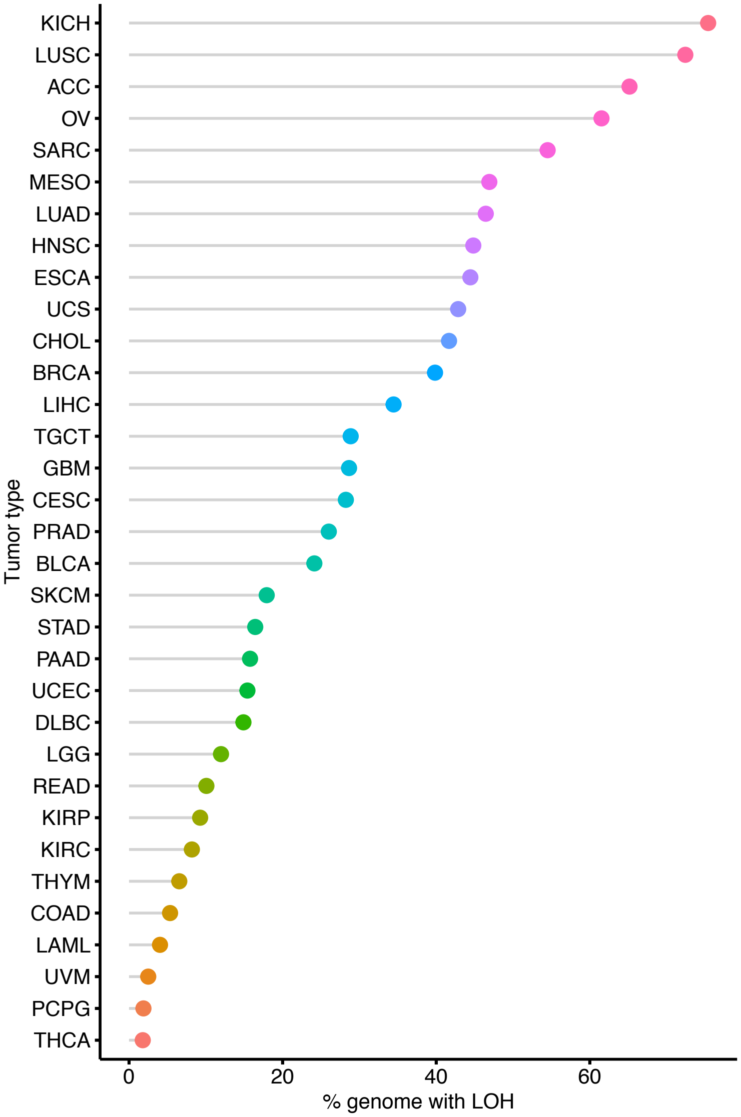


**Supplementary Figure 4.**
**a.** Table of *EXOSC8*rs117135638 SNP statistics, including calculation of theoretical number of treatable patients per year (see Methods). **b.** Cleveland dot plot of the rate of *EXOSC8* LOH across 33 TCGA tumor types. Tumor types are indicated by TCGA abbreviations (see https://gdc.cancer.gov/resources-tcga-users/tcga-code-tables/tcga-study-abbreviations).

**c**

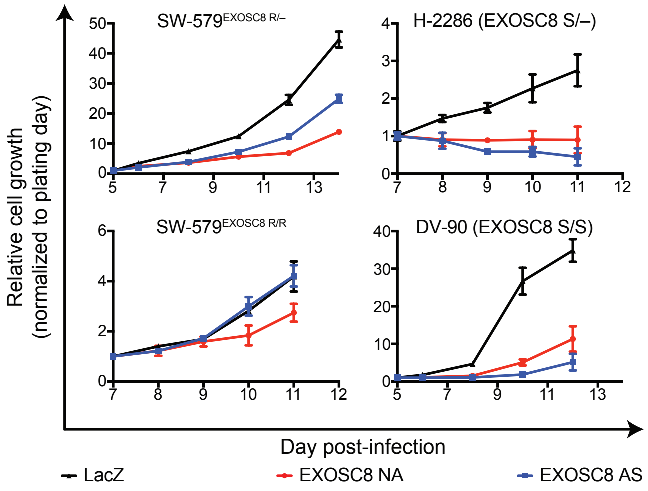


**d**

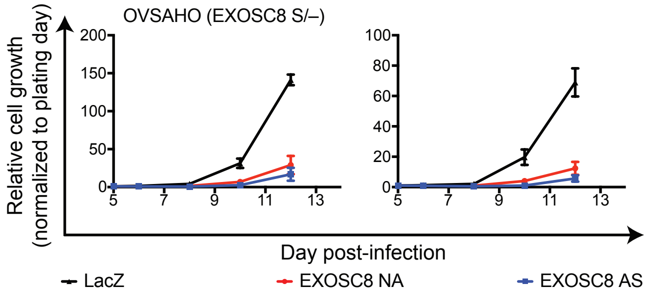


**e**

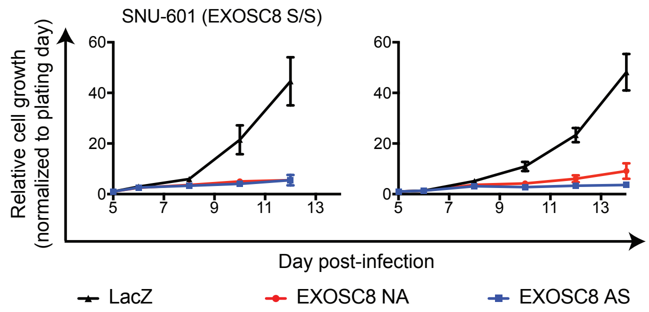


**f**

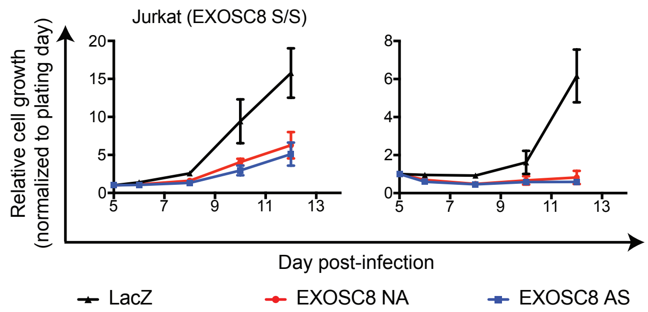


**Supplementary Figure 4.**
**c.–f.** Growth of indicated patient-derived cell lines expressing LacZ (black), EXOSC8 NA (red), or PRIM1 AS (blue) sgRNA as measured by CellTiter-Glo luminescence, relative to day of assay plating. n = 5 technical replicates; error bars represent s.d.

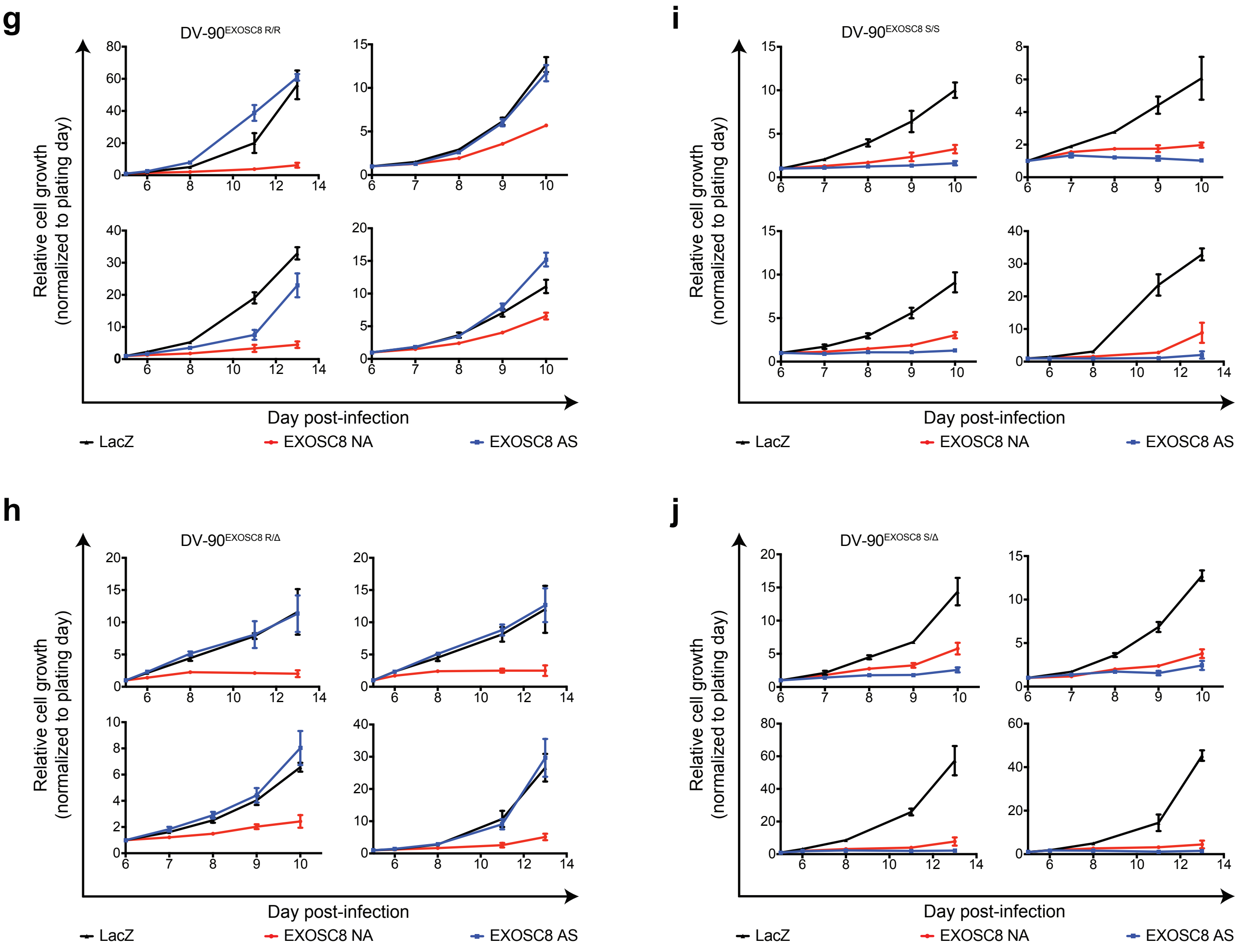


**Supplementary Figure 4.**

**g.–j.** Growth of indicated isogenic cell lines expressing LacZ (black), EXOSC8 NA (red), or PRIM1 AS (blue) sgRNA as measured by CellTiter-Glo luminescence, relative to day of assay plating. n = 5 technical replicates; error bars represent s.d.

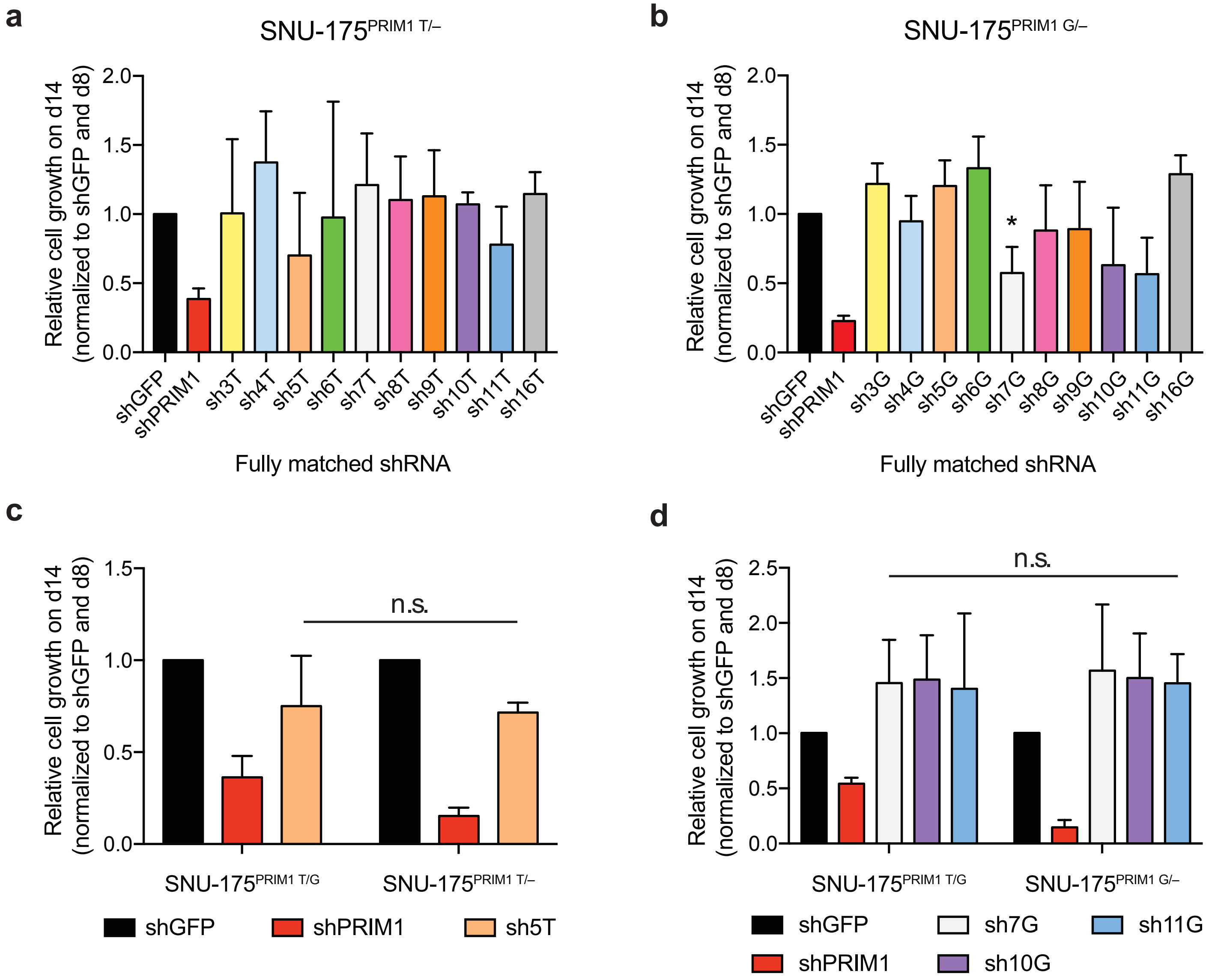


**Supplementary Figure 5.**
**a.** Growth of cells hemizygous for the major *PRIM1*$^{rs2277339}$ allele (SNU-175$^{PRIM1\ T/–}$) expressing shGFP, positive control shPRIM1, or indicated putative allele-specific shRNAs targeting the T allele of *PRIM1*$^{rs2277339}$, as measured by CellTiter-Glo luminescence, relative to day of assay plating. n = 3 biological replicates; error bars represent s.d. **b.** Growth of cells hemizygous for the minor *PRIM1*$^{rs2277339}$ allele (SNU-175$^{PRIM1\ G/–}$) expressing shGFP, positive control shPRIM1, or indicated putative allele-specific shRNAs targeting the G allele of *PRIM1*$^{rs2277339}$ as measured by CellTiter-Glo luminescence, relative to day of assay plating. n = 3 biological replicates; error bars represent s.d. One-tailed Student's t-test, *$p < 0.05$. **c.** Growth of heterozygous cells (SNU-175$^{PRIM1\ T/G}$) and cells hemizygous for the major *PRIM1*$^{rs2277339}$ allele (SNU-175$^{PRIM1\ T/–}$) expressing shGFP, positive control shPRIM1, or indicated putative allele-specific shRNAs targeting the T allele of *PRIM1*$^{rs2277339}$ as measured by CellTiter-Glo luminescence, relative to day of assay plating. n = 4 biological replicates; error bars represent s.d. One-tailed Student's t-test, n.s. = not significant. **d.** Growth of heterozygous cells (SNU-175$^{PRIM1\ T/G}$) and cells hemizygous for the minor *PRIM1*$^{rs2277339}$ allele (SNU-175$^{PRIM1\ G/–}$) expressing shGFP, positive control shPRIM1, or indicated putative allele-specific shRNAs targeting the G allele of *PRIM1*$^{rs2277339}$ as measured by CellTiter-Glo luminescence, relative to day of assay plating. n = 3 biological replicates; error bars represent s.d. One-tailed Student's t-test, n.s. = not significant.
